# Machine learning reveals bilateral distribution of somatic L1 insertions in human neurons and glia

**DOI:** 10.1101/660779

**Authors:** Xiaowei Zhu, Bo Zhou, Reenal Pattni, Kelly Gleason, Chunfeng Tan, Agnieszka Kalinowski, Steven Sloan, Anna-Sophie Fiston-Lavier, Jessica Mariani, Brain Somatic Mosaicism Network, Alexej Abyzov, Dimitri Petrov, Ben A. Barres, Hannes Vogel, John V. Moran, Flora M. Vaccarino, Carol A. Tamminga, Douglas F. Levinson, Alexander E. Urban

**Author notes:** Deceased, December 27, 2017 (Ben A. Barres). Full list of the members of the Brain Somatic Mosaicism Network is included after the acknowledgements.

## Abstract

Active retrotransposons in the human genome (L1, *Alu* and SVA elements) can create genomic mobile element insertions (MEIs) in both germline and somatic tissue^1^. Specific somatic MEIs have been detected at high levels in human cancers^2^, and at lower to medium levels in human brains^3^. Dysregulation of somatic retrotransposition in the human brain has been hypothesized to contribute to neuropsychiatric diseases^4,5^. However, individual somatic MEIs are present in small proportions of cells at a given anatomical location, and thus standard whole-genome sequencing (WGS) presents a difficult signal-to-noise problem, while single-cell approaches suffer from limited scalability and experimental artifacts introduced by enzymatic whole-genome amplification^6^. Previous studies produced widely differing estimates for the somatic retrotransposition rates in human brain^3,6–8^. Here, we present a highly precise machine learning method (RetroSom) to directly identify somatic L1 and *Alu* insertions in <1% cells from 200× deep WGS, which allows circumventing the restrictions of whole-genome amplification. Using RetroSom we confirmed a lower rate of retrotransposition for individual somatic L1 insertions in human neurons. We discovered that anatomical distribution of somatic L1 insertion is as widespread in glia as in neurons, and across both hemispheres of the brain, indicating retrotransposition occurs during early embryogenesis. We characterized two of the detected brain-specific L1 insertions in great detail in neurons and glia from a donor with schizophrenia. Both insertions are within introns of genes active in brain (*CNNM2*, *FRMD4A*) in regions with multiple genetic associations with neuropsychiatric disorders^9–11^. Gene expression was significantly reduced by both somatic insertions in a reporter assay. Our results provide novel insights into the potential for pathological effects of somatic retrotransposition in the human brain, now including the large glial fraction. RetroSom has broad applicability in all disease states where somatic retrotransposition is expected to play a role, such as autoimmune disorders and cancer.

## Introduction

Recent studies reporting on brain somatic MEIs have addressed the signal-to-noise problem to detect somatic MEIs either with a capture approach such as retrotransposon capture sequencing (RC-seq) from bulk brain tissue^12^, or with single-cell methods (because a somatic MEI is heterozygous within each mutated cell) including single-cell L1 insertion profiling (L1-IP)^13^, single-cell WGS (sc-WGS)^3^, and single-cell L1-associated variant sequencing (SLAV-seq)^8^. A limitation of these methods stems from the frequent occurrence of sequencing artifacts from chimeric DNA molecules resulting from the high numbers of PCR cycles (capture) or massive enzymatic whole-genome amplification (sc-WGS)^6^. It is also expensive to apply sc-WGS to, e.g., hundreds of cells in each of multiple brains and brain regions. An additional problem with any WGS approach is that MEI detection relies on uniquely mapping highly repetitive sequencing reads which remain difficult to resolve.

A less biased and more cost-effective approach is to identify somatic MEIs in deep ‘bulk-sequencing’ data: whole-genome sequencing (WGS) of genomic DNA extracted from bulk tissue, typically from over 10,000 cells (Fig. 1A, B). As in other methods^14–17^, MEI detection is then based on two types of sequencing reads (Fig. 1C): split-reads (SR), which capture the MEI insertion point such that part of the read maps to ME consensus sequence and the other part to the unique flanking reference sequence at the new genomic location; and paired-end (PE) reads where one read maps to ME consensus and the other to the unique flanking sequence. In either case, the unique sequence localizes the MEI in the genome. Existing algorithms based on these principles can detect germline MEIs^18^, or MEIs carried by a high subclonal fraction of tumor cells (>25%)^2^, but require many supporting reads per ME insertion for reliable detection. Lowering the detection threshold (number of supporting reads) leads to overwhelming numbers of false positives, likely from experimental noise and alignment errors. For example, using one supporting read in WGS data at 50× genomic coverage, we should detect ≥50% of MEIs that are present in ≥0.96% of cells. However, using a standard MEI algorithm ‘RetroSeq’^16^ to detect calls with one supporting read yielded ~59,900 (95% CI: 55,100-64,700) false positive MEI detections (Fig. 1D and Extended Data Fig. 1A).

**Fig. 1:**
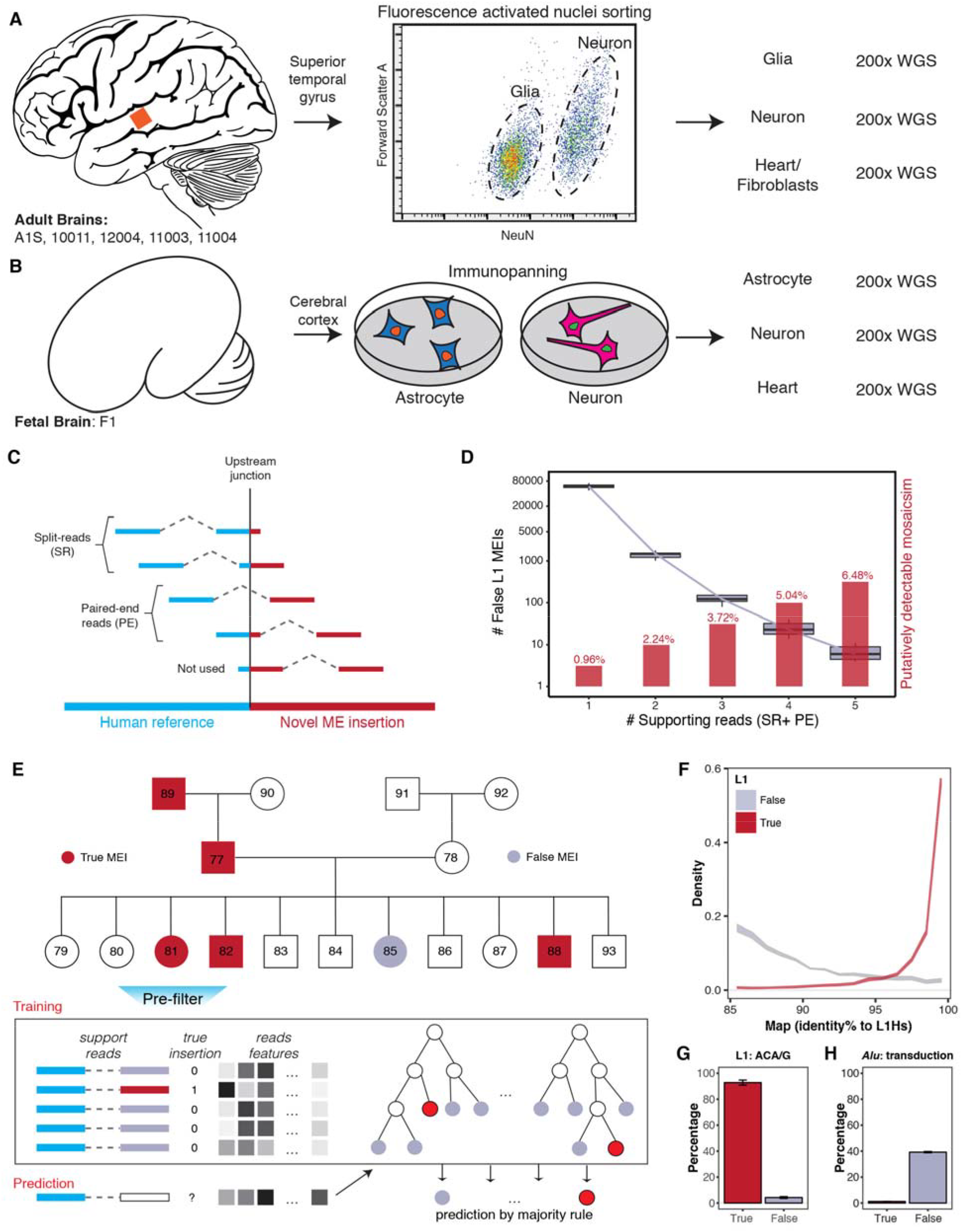
Project overview and machine learning method. (**A** and **B**) Deep genome sequencing of five adult brains and one fetal brain. For each donor, DNA from glia (astrocytes for “F1”), neurons, and a non-brain control tissue were sequenced to 200× genomic coverage. (**C**) Both split-reads (SR) and paired-end reads (PE) can be used to detect a mobile element insertion (MEI). Blue, segment of supporting read that maps to flanking sequence; red, segment of read that maps to ME consensus. (**D**) Detection of low-mosaicism MEIs requires a low-stringency for the number of supporting reads and is usually accompanied by many false positives. Gray, number of false positives vs. supporting-read cutoffs; red, theoretical lowest levels of detectable mosaicism vs. supporting-read cutoffs. (**E**) Training RetroSom. True (red) and false (gray) MEIs were labeled based on inheritance patterns, allowing for the training of a random-forest model using sequence features to classify supporting reads. A detailed flowchart of the modeling is shown in Extended Data Fig. 1B. (**F**) Supporting reads of true L1 MEIs (red) have a much higher homology to L1Hs consensus than reads supporting false insertions (gray). (**G**) True L1 supporting reads have the L1Hs-specific allele ACA/G. (**H**) True *Alu* retrotransposition does not include the flanking sequence from the putative source location. 95% confidence intervals are represented by the bandwidth (**F**) or error bars (**G, H**).

## Results

### Optimization of somatic MEI detection with machine learning

To better distinguish signal from noise in bulk sequencing, we developed RetroSom, which integrates RetroSeq (for mapping of reads to ME or reference sequence) with a transfer learning model trained on evolutionarily recent germline MEIs to detect low-level somatic MEIs. We trained RetroSom using polymorphic germline MEIs selected from Illumina Platinum Genomes WGS data^19^ for 17 members of a three-generation pedigree (Fig. 1E and Extended Data Fig. 1B). We assumed that recent germline MEIs would produce high-confidence non-reference calls that segregate in a Mendelian fashion. We excluded genomic regions with telomeric or centromeric repeats, segmental duplications, gaps, or near reference MEI insertions of the same type and on the same strand, totaling 21% of the genome for detection of *Alu* or 24% for L1. We also removed regions with abnormal sequencing depth, and supporting reads with low sequence complexity. We defined *true* MEIs based on inheritance pattern. Criteria for *false* MEI calls (likely artifacts) were <3 supporting reads in offspring and missing in both parents. We detected non-reference *true* insertions including, on average, 89 L1 and 467 *Alu* per offspring (Extended Data Fig. 1C). We then selected differentiating sequence features, ignoring the number of supporting reads (because this was a selection criteria for *true* MEIs) and features specific to individual elements (e.g., unique SNPs/Indels, unlikely to be shared by other families) or to sequencing conditions (to preserve generalizability) or chromosomal location -- positional bias could be due to natural selection or genetic drift and thus irrelevant to detecting somatic MEIs^20,21^. We chose 16-28 differentiating features each for four supporting-read classes (L1 and *Alu* PE and SR) (Supplemental Table 3).

We developed a machine learning algorithm using these features to classify *true* or *false* L1 or *Alu* supporting reads (Extended Data Fig. 1D, E). We tested logistic regression (with and without regularization), random forest and naïve Bayes classifiers, using 11× cross-validation (training on 10 offspring, testing on the eleventh). The random forest model performed best, with the area under the precision-recall curve at 0.965 (95% CI: 0.959-0.971) (Extended Data Fig. 1F, G). The most important differentiating features were sequence homology to the L1Hs or AluY consensus (Fig. 1F), L1Hs-specific SNPs (Fig. 1G)^22^, and exclusion of *Alu* calls with flanking sequence from the putative source location (“transduction,” which occurs with L1 but not *Alu* retrotransposition, Fig. 1H)^23^.

### Performance evaluation on three independent datasets

We tested RetroSom in several independent WGS datasets. Data from clonally expanded fetal brain cells^24^ confirmed that ≥2 supporting reads are necessary for high precision (L1: 99.97%; *Alu*: 99.99%) with adequate sensitivity (L1: 49.5%; *Alu*: 82.52%) (Fig. 2A and Supplementary Note 1). We also identified one somatic L1 insertion with features suggesting a rare non-TPRT, endonuclease-independent process (Supplementary Note 2)^25^. In addition, Illumina sequencing libraries prepared using a PCR-based method (~10 cycles) yielded 30-1000% more false MEIs than PCR-free libraries. RetroSom removed all false MEIs, yielding similar sensitivities between the two library types (L1: ~70%; *Alu*: ~86%) (Fig. 2B and Supplementary Note 3).

**Fig. 2:**
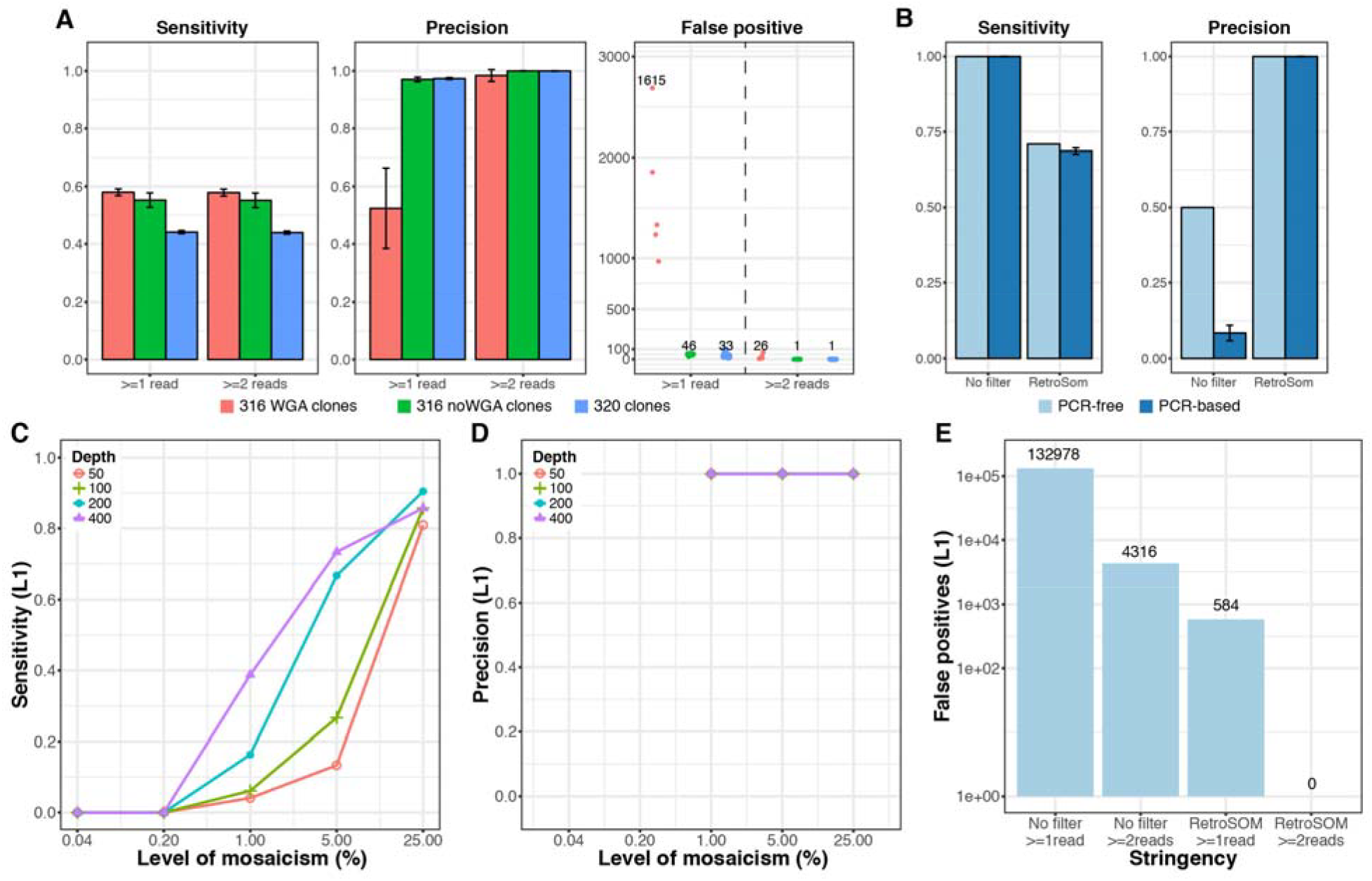
Benchmarking in independent test datasets. (**A**) Performance in detecting germline L1 insertions from clonally expanded fetal brain cells sequencing data. Red, clones from donor “316” sequenced with whole genome amplification (316WGA); green, the rest of the “316” datasets (316 noWGA); blue, clones from donor “320”. The error bars represent the 95% confidence intervals estimated in individual clones. (**B**) Performance in detecting germline L1 insertions from sequencing libraries prepared with or without PCR. A similar comparison for libraries created with a different tissue produced similar results (Extended Data Fig. 4). (**C-E**) Performance in detecting somatic MEIs simulated by six genomic DNA samples at proportions of 0.04% to 25% with that of NA12878, at various sequencing depth (red, 50×; green, 100× blue, 200× purple, 400×). Similar performance was observed for detecting *Alu* insertions (Extended Data Fig. 2).

We further benchmarked RetroSom with a genome mixing experiment. We pooled DNA from 6 human genomes (after calling high-confidence germline MEIs from available data) in precise proportions of 0.2%-25% with HapMap sample NA12878 (whose germline MEIs are also established); sequenced the pool (and NA12878 separately as a control) at 200× and called MEIs with RetroSom. A heterozygous germline MEI present in one of the six genomes will then occur as a mosaic MEI, with few or no supporting reads. RetroSom L1 detection sensitivities were 0 at mixing proportions of 0.04% and 0.2%, 0.16 at 1%, 0.67 at 5% and 0.90 at 25%, with no false positives (Fig. 2C, D). Detection rates were higher for RetroSeq alone (0.32 for 1%) or using RetroSom but requiring only one supporting read (0.48 for 1%), but with 4316 and 584 false positives, respectively (Fig. 2E). Sequencing depth, when computationally varied from 50×to 400×, linearly predicted detection sensitivity (especially for low proportions), but not precision (Fig. 2C-E). RetroSom was more sensitive and less precise for *Alu*, detecting 5 *Alu* at 0.2% mosaicism with 5 false positives (Extended Data Fig. 2C-E). Thus, using 200× WGS data, RetroSom can detect most mosaic L1 and *Alu* MEIs at >5% mosaicism, one-sixth with 1% mosaicism, and <1/100 with <0.2% mosaicism.

### Discovery and validation of somatic mobile element insertions

To detect real brain somatic MEIs, we separately analyzed neurons and glia (sorted by fluorescence-activated nuclear sorting) of five adult human postmortem brains: one elderly adult (“A1S”) and two schizophrenia-control pairs (Dallas Brain Collection)^26^, and neurons and astrocytes (sorted by immuno-panning) from one fetal brain (“F1”) (Supplementary Table 1). We collected superior temporal gyrus (STG) tissue from adult brains and cortical tissues from fetal brain, and heart or fibroblast control tissue, and sequenced extracted genomic DNA from each specimen to 200X whole-genome coverage (Fig. 1A, B). We called somatic MEIs supported with ≥2 high-confidence reads in either brain fraction but none in the corresponding control. As described above, we again excluded 24% of the genomic sequence from analysis for L1 and 21% for *Alu* MEIs. There were 0-3 somatic L1 and 0-13 somatic *Alu* calls per fraction (Supplementary Table 4). We selected MEIs for validation by blinded manual inspection with a novel visualization tool (RetroVis), following a checklist of selection criteria (Extended Data Fig. 5). We excluded most putative insertions, primarily those with potential mapping errors, and those identified in the 1000 genome project^27^ (likely germline insertions) (Supplementary Table 4). Two brain L1 insertions (L1#1, L1#2), both from the same donor with schizophrenia, met all criteria and were selected for in-depth investigation.

We validated both of these insertions following Brain Somatic Mosaicism consortium guidelines^28^. We quantitated mosaicism levels using droplet digital PCR (ddPCR), and determined the insertion sequences (single-base resolution) by nested PCR and Sanger sequencing. L1#1 was detected with two high-quality paired-end supporting reads in neurons, covering the upstream and downstream junctions (Extended Data Fig. 6A). Estimated mosaicism levels were 0.72% of neurons (95% CI: 0.50-0.94%), 0.54% of glia (95% CI: 0.40-0.67%) in the discovery region, and 0% in fibroblasts (8 technical replicates, Extended Data Fig. 6B, C). The full insertion sequence demonstrated four hallmarks of *in vivo* L1 retrotransposition (Extended Data Fig. 6D-F): (i) The endonuclease cleavage site is 5’-TTTT/CA-3’, similar to the degenerate consensus motif 5’-TTTT’AA-3’^29,30^. (ii) Consistent with the common 5’ truncation of new L1 insertions^31^, L1#1 is a 384bp 3’ fragment of L1 consensus, with a poly(A) tail of ~35bp (Extended Data Fig. 9E) and a short microhomology at the 5’ genomic DNA/L1 sequence junction, as preferred by 5’-truncated L1^32^. (iii) We confirmed a 15-bp target site duplication (TSD), as expected with TPRT retrotransposition. (iv) L1#1 carries the diagnostic ACA allele at base 5927-5929, the G allele at base 6012, and no other mismatches, indicating that the source element is from the youngest L1Hs-Ta subfamily (Extended Data Fig. 6D)^22^.

L1#2 was discovered with three supporting reads, including a split-read spanning the upstream junction (Extended Data Fig. 7). Estimated mosaicism levels were 1.2% of neurons [95% CI: 1.0-1.4%], 0.53% of glia [95% CI: 0.46-0.60%]), and 0% in fibroblasts (8 technical replicates, Extended Data Fig. 7B, C). The endonuclease site is 5’-CTTT/AA-3’, and the sequence contains a 418bp 3’ fragment of the consensus sequence, a poly(A) tail of ~25bp (Extended Data Fig. 9E) and a 6-bp TSD (Extended Data Fig. 7D). L1#2 also belongs to the L1Ta subfamily, with one mismatch with consensus sequence (Extended Data Fig. 7D).

### Spatial occurrence of somatic L1 retrotransposition in neurons and glia

Previous studies detected individual L1 insertions in neurons with narrow or broad distributions in one hemisphere^3^. Here, we quantitated the levels of L1#1 and L1#2 in neurons and glia from twenty-three brain regions from symmetrical sites of both hemispheres (Fig. 3 and Extended Data Fig. 9A). L1#1 was detected in neurons of all 23 regions (0.05-2.46% mosaicism), and in glia of 16 regions (0.05-3.46%) (Fig. 3A, C), including the putamen in the basal ganglia and the cerebellum, with the average of neurons and glia maximizing in left occipital cortex distal to STG (neurons, 0.95% (95% CI: 0.8-1.1%); glia, 3.5% (95% CI: 2.8-4.1%)). L1#2 was absent in specimens from prefrontal cortex, putamen and cerebellum. It was detected in 11 of 23 regions, all in the cerebral cortex (neurons: 0.1-1.4%; glia: 0.07-1.1%) (Fig. 3B, D), maximizing in right occipital cortex distal to STG. For both insertions, mosaicism levels were similar in neurons and glia from the same regions (Spearman *ρ*=0.77, *p*=5×10^−10^) (Extended Data Fig. 9F).

**Fig. 3:**
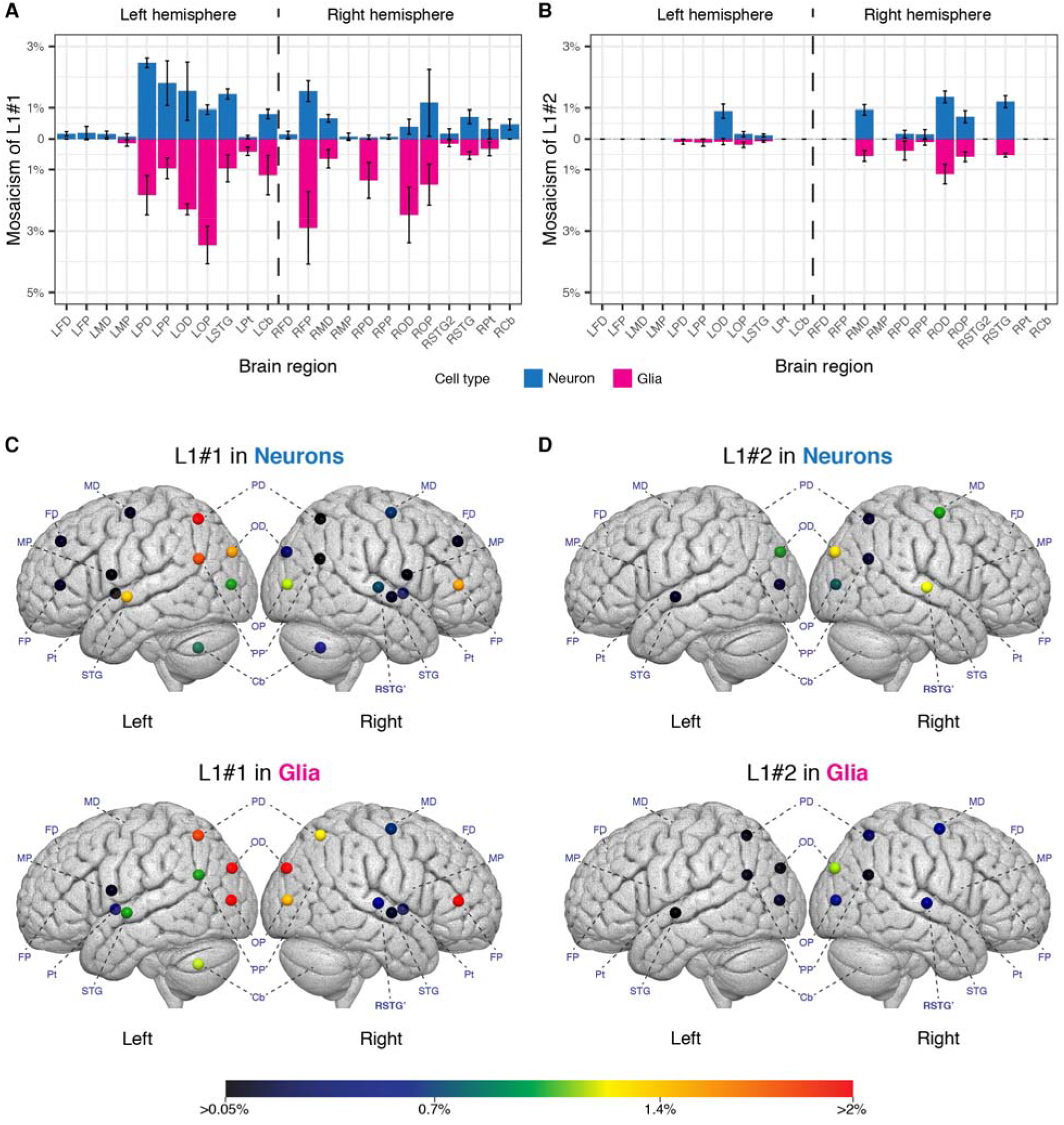
L1#1 and L1#2 have wide anatomical distribution in glia as well as in neurons. We quantitated the levels of mosaicism of two somatic L1 insertions, L1#1 and L1#2, in neurons and glia in 23 anatomical regions. (**A** and **B**) The levels of mosaicism and their 95% confidence intervals for L1#1 and L1#2 in neurons (blue) and glia (magenta). (**C** and **D**) Replotting the levels of mosaicism in the corresponding brain anatomical regions. L1#1 has a widespread pattern and is present in the neurons of all 23 brain regions, and the glia of 16 regions. L1#2 is present in 11 cerebral cortical regions. The level of mosaicism is denoted by a scale from cold (black, 0.05%) to hot (red, >2%). L, Left; R, Right; FD, prefrontal cortex – distal (to STG); FP, prefrontal cortex – proximal (to STG); MD, motor cortex – distal; MP, motor cortex – proximal; PD, parietal cortex – distal; PP, parietal cortex – proximal; OD, occipital cortex – distal; OP, occipital cortex – proximal; STG, superior temporal gyrus (2^nd^ sample); Pt, putamen; Cb, cerebellum; **RSTG’**, Right superior temporal gyrus (site of discovery). The exact anatomical locations are labeled in Extended Data Fig. 9A.

### Dysregulation of gene expression by L1 insertion

Both somatic L1 insertions occur in genomic regions of high functional potential. L1#1 is inserted in an intron of *CNNM2* (antisense strand), while L1#2 is in an intron of *FRMD4A* (sense strand). Specifically, L1#1 is inserted within a 1639bp putative transcriptional regulatory element ENSR00000032826 (Ensembl v95, Fig. 4A) that is defined by histone modification markers (Supplemental Table 6)^33^, in a broad linkage disequilibrium region surrounding *AS3MT* and *CNNM2* where genome-wide significant evidence for association was reported for schizophrenia^9^ and other traits (Fig. 4A, Extended Data Fig. 8, and Supplemental Table 7)^34^.

**Fig. 4:**
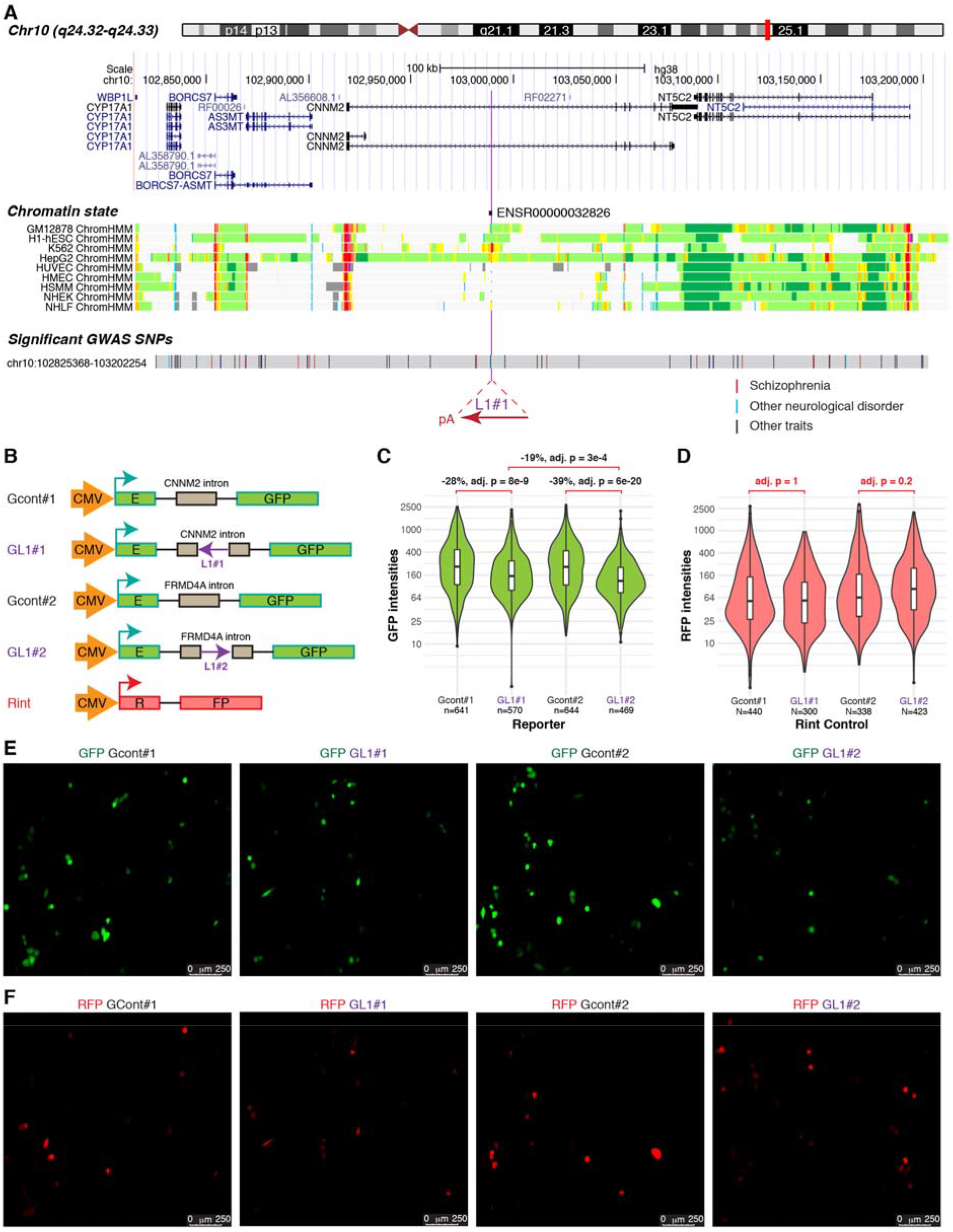
Intronic L1 insertions suppress EGFP reporter activities. (**A**) L1#1 is inserted in a 1639bp promoter flanking region (ENSR00000032826) that is expected to regulate the expression of nearby genes^33^. The chromatin states are shown for a subset of human cell lines: light gray, heterochromatin; light green, weakly transcribed; yellow, weak/poised enhancer; orange, strong enhancer; light red, weak promoter; bright red, strong promoter^38,39^. L1#1 is inserted in a linkage disequilibrium (LD) block, based on the common SNPs that are highly correlated (R^2^ > 0.6) with the closest common SNP to L1#1, rs1890185 (398bp upstream of L1#1). This LD block (gray) contains 72 SNPs significantly associated with 10 diseases or disorders and 28 measurement or other traits, including 13 risk SNPs from 11 schizophrenia studies^34^. Red, SNPs associated with schizophrenia; blue, SNPs associated with other neurological disorders; black, SNPs associated with other traits. (**B**) L1#1 and L1#2, as well as their flanking sequences, were cloned into a constitutively spliced intron in an EGFP reporter. An unmodified RFP reporter (Rint) was used as a control. (**C**) Cells transfected with either L1 insertion produced significantly less fluorescence than the controls in experiment (**B)**, and L1#2 has a stronger effect than L1#1. (**D**) The red fluorescence is generally consistent across assays, except for a slight increase in the cells transfected with L1#2. (**E**) A representative of the green fluorescence in cells transfected with L1 or control reporters. (**F**) A representative of the red fluorescence of the control plasmid ‘Rint’.

*CNNM2* and *FRMD4A* are expressed in many tissues, with higher levels in brain^35,36^. Tissue culture studies show that intronic L1 insertions, on either strand, can alter or disrupt gene expression, e.g., by inhibiting transcription elongation, altering splicing, terminating transcription prematurely or modifying local chromatin structure^37^. The effect depends on insertion length, strand, and splicing or polyadenylation sites within the insertion. Here we obtained the first experimental data showing the effects on gene expression of mosaic L1 insertions discovered in primary human brain cells.

Using a green fluorescent protein (EGFP) reporter “Gint”^40^, we tested the effects of L1#1 and L1#2 on gene expression by cloning each one (with flanking sequence) into a constitutively spliced intron in the antisense or sense strand respectively of the EGFP locus (Fig. 4B and Extended Data Fig. 9B). Control reporters were generated for the two flanking sequences without L1 insertion, with red fluorescent protein (RFP) reporter ‘Rint’ as an internal control^40^. In a blinded experiment, we transfected each of the modified Gint reporters with Rint control into HeLa cells and measured the level of fluorescence (Fig. 4C-F). Compared to controls, L1#1 (antisense) reduced green fluorescence by 28% (95% CI: 20-35%, *t*=-6.2, adjusted *p*=8×10^−9^), and L1#2 (sense) reduced green fluorescence by 39% (95% CI: 33-45%, *t*=-9.6, adjusted *p*=6×10^−20^) (Fig. 4C). Including the intronic length as a covariate, the difference in fluorescence remains significantly correlated for insertion vs. control assay (*t*=-9.27, adjusted *p*=4×10^−19^). There was also a modestly significant difference in reductions of green fluorescence for L1#1 vs. L1#2 insertions (*t*=-4.12, adjusted *p*=3×10^−4^), possibly due to a weak polyadenylation signal in the L1#2 sense strand (but not the truncated L1#1 sequence) that could terminate transcription prematurely^37^. The red fluorescence was generally consistent across all assays, except for a slight increase in assay L1#2 *(t*=2.4, adjusted *p*=0.2), possibly due to weaker competition from EGFP synthesis in the same cells (Fig. 4D). We confirmed similar results in a separate experiment with no Rint control (Extended Data Fig. 9C, D). These *in vitro* results suggest that L1#1 and L1#2 could, in principle, reduce expression of genes into which they are inserted.

## Discussion

We demonstrate that each of the two somatic MEIs that we characterized in greater detail has similar mosaicism levels and anatomical distributions for glia and neurons. Thus glia, which are at least equal in numbers to neurons amongst the cells of the human brain, are similarly important targets for research on the physiological impact of somatic retrotransposition and the tracing of developmental lineages. The distributions of the two validated L1 insertions in both neurons and glia (but not fibroblasts) indicate that the retrotransposition likely occurred in neuroepithelial cells at the neural plate stage: before the separation of cerebellum, basal ganglia and cortex for L1#1, and later in a dorsal telencephalic neuroepithelial cell for L1#2. Both neuroepithelial cells would give rise to bipotential neural stem cells (the radial glia)^41^, that develop into neurons and glia and serve as a guiding scaffold for their migration from the developing ventricular zones to the cortical surface. Thus L1#1 and L1#2 both have similar mosaicism levels in neurons and glia from the same anatomical region, with the earlier mutation event (L1#1) producing higher mosaicism levels.

Widely divergent estimates of the rate of somatic L1 insertions in brain have been reported. Two previous unsorted bulk sequencing studies described hundreds of putative somatic L1 insertions at 80× coverage using Complete Genomics sequencing^12^ or thousands per region at 30× coverage using targeted Illumina sequencing^5^. Our mixing experiment results suggest that sequencing at these depths would only detect insertions with higher mosaicism levels than were observed here: our sensitivity to detect mosaicism levels >5% was 0.67, and none were observed, suggesting that such events are rare. Our data are more consistent with careful analyses of single-cell data, with extensive computational filtering and experimental validation, which estimated <0.2 L1 retrotransposition events/neuron^3,6,8,13^. RetroSom’s sensitivity is ~0.16 for insertions at 1% mosaicism (Fig. 2C), but it masks 24% of the genome as unreliable for detection. Assuming new L1 retrotransposition occurs at a similar rate in highly repetitive regions and in reference genomic L1s^21,42^, average sensitivity is therefore reduced to ~0.12, thus ~1/8th of MEIs at 1% mosaicism are detected. We validated two somatic L1s from the brain selected for extensive characterization, at 0.72-1.2% mosaicism at the discovery site, which should therefore contain 8 x 2 = 16 somatic L1 insertions at ~1% mosaicism levels. Thus, sampling single cells in this region would yield on average 0.16 somatic L1 insertions per cell, similar to the previous single-cell estimates.

Future developments in WGS will accelerate this area of research. Using even deeper WGS coverage will allow for detection of very low (<<1%) mosaicism levels, e.g., in fetal brain tissues where full clonal expansion has not occurred, or after sorting for highly differentiated cells with limited further expansion, or regions where mosaicism level is modified by the degree of tangential migration and neurodevelopmental programmed cell death^43^. Accurate and cost-effective long-read WGS technologies will emerge and allow to improve somatic MEI detection in repetitive sequences, and longer reads or longer paired-end insert sizes will permit detecting mosaic L1s carrying 5’ or 3’ sequences from the source region (transduction)^23^. The precision of our modeling strategy is constant with higher depth and read length, while sensitivity will improve.

Can moderate or low levels of mosaicism have pathological consequences? Known disease-associated MEIs are *germline* mutations^44^. In human brain, somatic single nucleotide variants (SNVs) with low tissue allele frequencies (tAF, the fraction of chromosomes carrying an alternative allele) in known risk genes can drive functional anomalies^28^, such as Sturge-Weber syndrome (1-18% tAF)^45^, focal cortical dysplasia (1.3-12%)^46^, and hemimegalencephaly (8-40% tAF)^47,48^. Our observation of somatic MEIs in 0.05-3.46% of brain cells in multiple regions translates to 0.025-1.73% tAF. Regarding whether such levels can be pathological, e.g., in neuropsychiatric disorders such as schizophrenia, one can consider a hypothetical scenario of an etiologically-relevant anatomical region where 3% of cells carry functionally damaging MEIs, i.e., acting as somatic additional genetic hits, potentially in an individual with a genomic background that already confers elevated polygenic disease risk. Taking our findings as examples, L1#1 disrupted *CNNM2*, in which common SNPs are strongly associated with schizophrenia (Fig. 4A)^9^ and which is likely a driver gene^10^; while L1#2 disrupted *FRMD4A*, associated with a syndrome of microcephaly and intellectual disability^11^, phenotypes observed in carriers of copy number variants that increase risk of schizophrenia^49^. Both genes are expressed in brain and both MEIs reduced gene expression in an experimental assay. L1#1 also disrupted a putative transcriptional regulatory element with predicted effects on multiple neighboring genes^50^, which could lead to rewiring of transcription networks.

These observations in one patient do not establish an etiological role. However, it is likely that each patient with a common genetically complex disease has a unique set of common variants with small effects on risk, and rare variants with larger effects^9^. The latter could include mosaic MEIs with strong local effects that extend beyond the mutated cells and thus are less dependent on mosaicism levels, such as disordered neurodevelopment, induction of epileptiform activity, disruption of brain circuit activity through the large numbers of synaptic connections of the mutated cells, or cell-cell contacts during epithelial cell polarization (e.g., the essential role played by *FRMD4A* in the cell adhesion protein complex)^51^. In diseases like schizophrenia or autism with greatly reduced fecundity, large-effect *de novo* mutations in germline and soma may play a role in maintaining disease frequency^52^. Thus, it is worth keeping an open mind about the potential pathological consequences of even low mosaic levels of somatic MEIs in neuropsychiatric disorders. Finally, while high-frequency somatic MEIs are described in human cancers^2^, lower-frequency somatic mutations can also be clinically significant, due to tumor impurity and subclonality^53^. RetroSom could be applied to investigate the full frequency spectrum and clonal architecture of MEI mosaicism in tumors, as well as other disease states where somatic retrotransposition is expected to play a role.

## Notes

#### Summary of Updates

Changed the subject area from “bioinformatics” to “genomics” Separated the manuscript into “main text” and “supplementary materials”

