## Supplementary Materials for "Machine learning reveals bilateral distribution of somatic L1 insertions in human neurons and glia"

METHODS

**Whole genome sequencing of six human donors**

We studied 6 human donors in this project, including an adult donor A1S, a fetal donor F1, and two schizophrenia-control pairs matched as closely as possible for age, brain pH, and postmortem delay to autopsy: “10011”, “11003”, “11004”, and “12004” (Supplementary Table 1). We obtained postmortem brain tissue and heart tissue for donors A1S and F1 with informed consent under a Stanford University Institutional Review Board approved protocol. Human brain tissue and fibroblast from the schizophrenia and control donors were obtained from the Dallas Brain Collection^1^. The clinical diagnosis for each of the schizophrenia/control donors was evaluated by at least two research psychiatrists. The schizophrenia/control status was initially masked during MEI discovery.

**Fluorescence-activated Nuclear Sorting (FANS)**

For the initial whole genome sequencing screening of the adult donors, we sampled 0.5-1 cm^3^ cortical tissues from the superior temporal gyrus (STG). The neuronal and glial nuclei were extracted from the postmortem brains using methods modified from a published protocol^2^. Briefly, the brain tissues were dissected on a cold plate (TECA™ LHP-1200CAS) into ~200mg segments. For each segment, we homogenized the tissue in 3.6ml lysis buffer (0.32M sucrose, 5mM calcium chloride, 3mM magnesium acetate, 0.1mM EDTA, 1mM DTT, 0.1% TritonX-100, and 10mM Tris PH 8.0). We then added 6.5ml sucrose buffer (1.8M sucrose, 3mM magnesium acetate, 1mM DTT and 10mM Tris PH 8.0) to the bottom of the tissue lysate, and centrifugated at 100,000g for 2 hours at 4 °C (Sorvall™ ultracentrifuge WX-80). The nuclei in the pellet were collected by incubation in 500 μl ice-cold PBS for 10 min, gentle resuspension, and filtration through a 40 μm strainer. We stained the nuclei with an anti-NeuN-PE antibody (Milli-Mark FCMAB317PE, 1:100), 1mg/ml DAPI (1:1000), and 10%BSA (1:50) for 45 min at 4 °C. The labeled nuclei were evaluated under a fluorescent microscope (EVOS FL), and the yield was quantitated with a hemocytometer.

The neuronal and glial nuclei were separated with fluorescence-activated nuclear sorting (FANS) using a BD Aira sorter that was optimized to sort nuclei based on DAPI and PE signals^3^. We first drew gates in forward scatter (FSC-A and FSC-W), side scatter (SSC-A and SSC-W), and DAPI channels to select for singlet nuclei. The NeuN+ and NeuN- nuclei were then separately collected with gates in the PE and FSC-A channels: NeuN+ nuclei are from neurons and are larger in size and carry stronger PE signals, while NeuN- nuclei are from non-neurons (glial cells) and are smaller. The purity of the sorted nuclei (quantitated by reanalyzing the sorted fractions) was >99.95% in both fractions. The data were analyzed with FlowJo cell analysis software (v10.0.7.r2). A typical yield from 200mg of brain tissue is 1-2 million nuclei, NeuN+ and NeuN- combined. The ratio between the NeuN+ and NeuN- fraction varies depending on the anatomical region, e.g.,1.6 in superior temporal gyrus, 12.6 in cerebellum, and 0.24 in putamen.

**Immuno-panning**

Immuno-panning was performed using methods modified from a published protocol^4^. In brief, fetal cortex was harvested from the elective termination of a gestational week 18 pregnancy. Cortical tissue was chopped into fine pieces (<1 mm^3^) with a #10 scalpel blade and then incubated in 15 U/mL papain at 34°C for 60 minutes. After digestion, the tissue was washed with a protease inhibitor stock solution. The tissue was then gently triturated to yield a single-cell suspension, which was added to a series of plastic petri dishes pre-coated with cell-type-specific antibodies. The antibodies used included anti-CD45 (BD 550539) to capture myeloid cells, anti-HepaCAM (R&D MAB4108) to capture astrocytes, anti-Thy1 (BD 550402) to capture neurons, and O4 hybridoma for oligodendrocyte lineage cells. The general scheme for isolating cell populations involved negative selection of ‘contaminating’ cell populations, followed by positive selection of the cell type of interest. For neurons, we first negatively selected contaminating cell types by immunopanning with anti-CD45, followed by two sequential anti-HepaCAM plates to deplete myeloid cells and astrocytes, respectively. The remaining cell suspension was then immunopanned with anti-Thy1 to positively select for fetal neurons. The general scheme for isolating astrocytes involved negative immunopanning with anti-CD45, followed by two sequential anti-Thy1 plates and two sequential anti-O4 plates to deplete myeloid cells, neurons, and oligodendrocytes, respectively. The remaining cell suspension was then immunopanned with anti-HepaCAM to positively select for fetal astrocytes. Cells were incubated on each immunopanning dish for 10-20 minutes at room temperature. Unbound cells were transferred to the subsequent petri dish, and the dish with bound cells was rinsed with PBS to wash away loosely attached contaminants. Adherent cells were dislodged with Trypsin (200 units in EBSS for 5 min at 37°C), which was briefly inactivated with FBS before spinning and resuspending purified cells.

**Genomic DNA extraction and whole genome sequencing**

The genomic DNA from neuronal nuclei, glial nuclei, and non-brain controls were extracted with the Qiagen DNeasy Blood & Tissue Kit. The yield is typically ~3 μg per million cells, and all DNA quality passed a DNA integrity number (DIN) threshold of 7. We prepared six separate libraries for each DNA specimen, using 200 ng genomic DNA and the Illumina TruSeq Nano DNA Sample Preparation Kit (Macrogen). These libraries were sequenced to >30x on an Illumina HiSeq X system, with a read length of 2x150 bp. For comparison, we also prepared two PCR-free libraries from A1S heart and A1S neuronal nuclei, each using 1 μg genomic DNA and the Illumina TruSeq DNA PCR-free Sample Preparation Kit.

RetroSom pipeline

Additional public datasets

We obtained several high-quality public whole genome sequencing datasets ­­for the training and testing of RetroSom (Supplementary Table 2), including:

a. Illumina Platinum Genomes

The Illumina Platinum Genomes dataset includes the CEPH pedigree 1463, with 4 grandparents (NA12889, NA12890, NA12891 and NA12892), 2 parents (NA12877 and NA12878), and 11 offspring (NA12879, NA12880, NA12881, NA12882, NA12883, NA12884, NA12885, NA12886, NA12887, NA12888 and NA12893)^5^. All members were sequenced to an average depth of 50x (dbGAP accession: phs001224). In addition, NA12877 and NA12878 were sequenced to an average depth of 200x (ENA accession: PRJEB3246). The sequencing was carried out in PCR-free libraries on an Illumina HiSeq 2000 system, with a read length of 2x101 bp.

b. Human Genome Structural Variation Consortium

We used whole genome sequencing data from three trios studied in the Human Genome Structural Variation (HGSV) Consortium, including Lymphoblastoid cell lines of a Yoruban trio (NA19238, NA19239 and NA19240), a Puerto Rican trio (HG00731, HG00732, and HG00733), and a southern Han Chinese trio (HG00512, HG00513 and HG00514)^6^. Each cell line was sequenced with PCR-free libraries to an average depth of >30x (ftp://ftp.1000genomes.ebi.ac.uk/vol1/ftp/data_collections/hgsv_sv_discovery/data/).

c. Clone sequencing datasets

The clone sequencing datasets 316 and 320 were downloaded from the NIH National Institute of Mental Health (NIMH) Data Archive (https://data-archive.nimh.nih.gov) under collection ID #2330 and DOI:10.15154/1410419^7^. Both datasets include whole genome sequencing of cell clones expanded from individual neural stem cells. Dataset 316 has 5 clones amplified with multiple displacement amplification (316WGA, N=5), along with 8 other clones and bulk DNA from the frontal lobe and spleen (316noWGA, N=10); dataset 320 contains 50 clones plus bulk DNA from the basal ganglia, frontal lobe, and spleen (320, N=53).

d. Brain Somatic Mosaicism Network (BSMN) Consortium common brain

We also obtained the sequencing data of the common brain tissue studied by the BSMN Consortium. The data include >200x whole genome sequencing of the bulk brain tissue and fibroblast.

Sequence alignment and candidate supporting reads

Raw sequencing reads from the six human donors, as well as from the public datasets, were all aligned to the human reference genome GRCh38DH with the Burrows-Wheeler Aligner (BWA v0.7.12; ‘mem -t 6 -B 4 -O 6 -E 1 -M -R’), and then postprocessed on alternative contigs/decoy/HLA genes (bwa-postalt.js)^8^. The alignment was further cleaned by removing secondary alignment, supplementary alignment, and PCR duplicates. We used a modified Retroseq pipeline^9^ (-discover -align -srmode -minclip 20 -len 26) to extract candidate supporting reads with >80% identity matching the consensus sequences of L1Hs or AluY elements, including AluYa5, AluYa5a2, AluYb8, AluYb9, AluYc1, and AluYk13^10^. We inferred MEIs by integrating two types of supporting reads: split-reads (SR), which capture the MEI insertion point such that part of the read maps to the ME consensus sequence and the other part to the unique flanking reference sequence at the new genomic location; and paired-end (PE) reads where one read maps to the ME consensus (ME end) and the other to the unique flanking sequence (anchor end). The two paired-end supporting reads are not properly paired because the ME end is usually mapped to a distant reference ME, and the sequence between the two paired reads is unknown but has a known size range. Thus, PE supporting reads help to localize the MEI without giving information regarding the exact breakpoints. The SR supporting reads, on the other hand, provide breakpoint sequences but are not always available when the insertion is found in a minority of cells.

The SR supporting read has one chimeric read mapped to both the flanking sequence and the ME sequence, and often contains too few base pairs of the flanking sequence for correct mapping. Thus, the correct placement of a chimeric read requires the mate-read to be properly paired. However, BWA-MEM sometimes assigns an incorrect primary alignment location for the chimeric read even when it is properly paired with its mate. BWA assigns two alignments for each chimeric read: a primary alignment based on the longer segment and a supplementary alignment based on the shorter segment. When a chimeric read covers a MEI junction, either segment can be in the flanking sequence and properly paired with the mate, while the other segment will be in the ME sequence and usually mapped to a distant reference ME. When the ME segment is >50% of the chimeric read in a SR supporting read, the chimeric read is mapped to a location not properly paired with its mate in the primary alignment. As a result, the supporting read will be reported as PE instead of SR, and the insertion junction information is lost.

To optimize the discovery of SR supporting reads, we scanned the supplementary alignment (SA tag) of all of the PE supporting reads for chimeric alignments. If the position of the shorter segment could be properly paired with the anchor end, and the longer fragment could be mapped to a ME sequence, we converted the PE supporting reads to SR. Furthermore, we separately analyzed a group of PE supporting reads with a split-read anchor end: the chimeric anchor end also provides vital information about the MEI junction. We ignored the PE supporting reads when <50% of their anchor ends were mapped to the flanking sequence, to avoid potential mapping errors.

We excluded supporting reads of poor quality, including those characterized by (1) genomic regions of highly repetitive sequences, including centromeric repeats, telomeric repeats, large segmental duplications, reference genome gaps, or within 100bp of a reference MEI of the same type and strand; (2) supporting reads with low sequencing complexity (SEG < 1)^11^; or (3) outlier sequencing depth within 500bp upstream and downstream to the insertion (>3 standard deviations away from the mean). The sequencing depth for sex chromosomes was evaluated separately. The masked reference sequence was 23.6% for L1 insertions in the positive strand, 23.7% for L1 insertions in the negative strand, 21.0% for *Alu* insertions in the positive strand, and 21.1% for *Alu* insertions in the negative strand.

Simulating the putatively detectable mosaicism

We performed a simulation to evaluate the relationship between the sequencing depth, number of supporting reads, and the detectable mosaicism of somatic MEIs (Extended Data Fig. 1A). In the simulation, we assumed that (i) sequencing depth is 50x; (ii) sequencing reads are 2x150bp in length and the fragment length (including read1, read2, and the insert in between) follows a normal distribution: $\mathcal{N}\left( 600, 100 \right)$; (iii) the MEI is from 4500bp to 5500bp on a DNA segment that is 10kb long; (iv) the MEI has no transduction; (v) the MEI is heterozygous in the somatic cells; (vi) the sequencing fragment is shorter than the MEI and thus cannot span both upstream and downstream junctions; (vii) any reads that cross the MEI junction with >30bp overlapping with the ME consensus and >half of the read length (75bp) overlapping with the flanking sequences can be used as supporting reads (i.e., the flanking sequence can be uniquely mapped); (vi) there are no split-read supporting reads from the MEI junction around the poly(A) tail because the poly(A) tail may cause inaccurate mapping of the split-read (Extended Data Fig. 4).

Under these assumptions, we define the *putatively detectable mosaicism* as the lowest mosaicism at which ≥50% of MEIs can be detected with a certain number of supporting reads. For instance in a hypothetical 50x WGS dataset, the 10kb DNA fragment containing the MEI in 0.96% of cells is expected to be covered with 8 read-pairs, and 52% of these MEIs are detectable with 1 or more supporting reads in 50000 simulations. Similarly, the *putatively detectable mosaicism* is 2.24% for 2 supporting reads, 3.72% for 3 supporting reads, 5.04% for 4 supporting reads, and 6.48% for 5 supporting reads (Fig. 1D). The real *detectable mosaicism* is likely higher because MEI supporting reads have to meet additional criteria, such as unique and high quality mapping of the anchor-end reads. The code for the simulation is available at https://github.com/XiaoweiZhuJJ/RetroSom.

Model training

We built the RetroSom model to classify each supporting read identified in the 11 offspring from the platinum pedigree as either a *true* or *false* MEI (Extended Data Fig. 1B). For all members in the pedigree, we first identified candidate MEIs with ≥1 support reads after excluding reference MEIs, regions of highly repetitive sequences, low sequencing complexity, or outlying read depth. Notably, we also separated the supporting reads from different DNA strands and called MEIs in forward/reverse strands separately. We then labeled each candidate MEI in the 11 offspring as *true* or *false* insertions based on the inheritance pattern. *True* insertions were transmitted from heterozygous or homozygous insertions in the parents (NA12877/NA12878). A heterozygous MEI satisfies three conditions: (1) found in a total of 1-10 offspring, each with >4 supporting reads; (2) found in NA12877 or NA12878, but not both, with >4 supporting reads; and (3) found in at least one of the two grandparents from either the maternal or the paternal side, but not both sides, with >4 supporting reads. A homozygous MEI satisfies another set of three conditions: (1) found in all 11 offspring with >4 supporting reads; (2) found in NA12877 or NA12878, but not both, with >4 supporting reads; and (3) found in both grandparents on either the maternal or the paternal side, but not both sides, with >4 supporting reads. We excluded MEIs present in both parents to remove common artifacts and evolutionarily-ancient insertions. As expected, the occurrence of *true* MEIs in offspring follows a binomial distribution (Extended Data Fig. 1C). The *false* insertions, on the other hand, are the ones found in the offspring but absent in both parents. There are substantial numbers of *false* insertions at a low cutoff of supporting reads (Fig. 1D**)**. In the *false* dataset for training, we only kept low confidence MEIs (<3 supporting reads) that are absent in both parents to exclude true *de novo* germline insertions in the offspring.

We built a data matrix with “positive” supporting reads from *true* MEIs and “negative” supporting reads from *false* MEIs; each read is characterized by a list of sequencing features (Supplementary Table 3). We built separate random forest models for L1 PE reads, L1 SR reads, *Alu* PE reads, and *Alu* SR reads to separate the positives from the negatives, using the selected sequencing features. The machine learning was carried out in R (v3.5.0): Missing values are known to cause problems in a random forest model. Thus, we partitioned L1 PE reads into 8 subgroups, with reads mapped to different segments of the L1 consensus (Extended Data Fig. 1D); L1 SR reads into 2 subgroups, including the original SR reads and the ones converted from PE reads; and *Alu* PE reads into 2 subgroups, including the ones with and without split-read anchor ends.

When applying the sub-models to make new predictions, one candidate L1 PE supporting read may be categorized to several subgroups and therefore have multiple probability scores. RetroSom reports the probability based on the submodel with the best accuracy, in the following order: (1) RFI.1, (2) RFI.4, (3) RFI.8, (4) RFI.2, (5) RFI.5, (6) RFI.7, (7) RFI.6, (8) RFI.3. The order is based on the overall accuracy of each model in the training dataset (Extended Data Fig. 1E). Most sub-models produced highly similar predictions and the ranking had little impact on the overall prediction. We chose the default probability score cutoff (>0.5) for classifying new supporting reads as true MEI insertions.

Evaluation training data with 11x cross validation

The performance of RetroSom was first evaluated with 11x cross validation. Each of the 11 offspring was selected as the test dataset once, while the data from the remaining 10 offspring were used for modeling. For comparison, we also built a logistic regression model (LogR), a Lasso regression model (Lasso), a Ridge regression model (Ridge), and a Naïve Bayes model. ­The machine learning was carried out in R (v3.5.0): logistic regression (with and without regularization)­ used the ‘glmnet’ package (v2.0-16); random forest used the ‘randomForest’ package (v4.6-14); and naïve Bayes used the ‘e1071’ package (v1.6-8)^12,13^.

We evaluated the models using six metrics: $accuracy= \left( TP+TN \right)/\left( TP+FP+TN+FN \right)$,$F_{1}={2TP}/\left( 2TP+FP+FN \right)$, $sensitivity={TP}/\left( TP+FN \right)$, $precision={TP}/\left( TP+FP \right)$, area under receiver operating characteristic curve (AUROC), and area under precision-recall curve (AUPR). $TP$, true positive; $TN$, true negative; $FP$, false positive; $FN$, false negative. AUROC and AUPR were calculated with the ‘PRROC’ package (v1.3.1) (Extended Data Fig. 1F, G)^14^.

Evaluation in fetal brain clonal expansion

We evaluated RetroSom in two public clone sequencing datasets, 316 and 320, created by culturing individual neural cells from fetal brains and sequencing genomic DNA from each clone ^7^. Dataset 316 includes 13 clones, 5 using whole genome amplification (WGA), and bulk brain and non-brain tissue; dataset 320 contains 50 clones and bulk DNA from two brain regions and one non-brain tissue. In addition to being single-cell clones, these datasets differed from the Platinum dataset in sequencing method (150bp reads vs. Platinum’s 101bp reads); use of WGA in 5 of the clones for 316 (analyzed separately); and lack of family data to define true MEIs. *True* MEIs in clonal data were defined as those supported in most clones (>4 supporting reads in >80% of clones) and *false* MEIs as insertions with <3 supporting reads in >80% clones. MEIs that have many supporting reads in individual clones but are missing in others could be *true* *de novo* insertions, and thus were excluded from both the *true* and *false* groups.

Evaluation in PCR-free sequencing libraries

We re-sequenced two specimens, A1S heart and A1S NeuN+, to 30x-coverage, using PCR-free sequencing libraries and 1 μg of genomic DNA each, and compared the MEI calling accuracy to two sets of six PCR-based (TruSeq Nano, ~10 PCR cycles) datasets created from the same tissues (Extended Data Fig. 4). The *true* and *false* MEIs of A1S were selected based on their presence in all 20 libraries, including 18 TruSeq Nano (3 cell fractions) and 2 PCR-free sequencing datasets. *True* MEIs were selected as the insertions that were highly supported in most of the libraries (>4 supporting reads in >80% libraries), while *false* MEIs were selected as the insertions that were missing or poorly supported in most of the libraries (<3 supporting reads in >80% libraries).

Evaluation in mixed DNA with different frequencies

To evaluate RetroSom’s performance for detecting MEIs with low levels of mosaicism, we designed a sequencing experiment to use genomic DNA mixed at various frequencies to simulate real mosaic MEIs. We first spiked six unrelated genomic DNA in NA12878 DNA at a gradient of concentrations, including 1) A1S heart at 0.04%, 2) NA19240 at 0.2%, 3) HG00733 at 1%, 4) HG00514 at 1%, 5) BSMN common brain at 5%, and 6) NA12877 at 25%. The mixed DNA was meant to simulate a specimen carrying somatic MEIs of different frequencies, while pure NA12878 was meant to simulate a control specimen without any somatic MEIs. The DNA we spiked in was chosen based on three criteria. (i) The chosen DNA was either sequenced deeply (>200x) by our group (A1S heart and BSMN brain) or included as the child in trios chosen by the HGSV (NA19240, HG00733, and HG00514) or Platinum Genomes (NA12877 and NA12878). Based on the existing sequencing data, we created a high confidence catalogue of MEIs that are unique to each DNA. Notably, homozygous MEIs are presented in the mixed DNA at a frequency twice as high as the heterozygous MEIs. To better simulate real somatic MEIs that are almost certainly heterozygous when occurring, we only considered heterozygous MEIs in each of the spiked genomes. (ii) we chose DNA of distinct ancestries to maximize the number of unique MEIs at each mosaic level. Most of the genomic DNA has a low level of heterozygous L1 insertions that are not shared with anyone else (between 11 and 32), except for the African sample NA19240, which has 77 unique L1. We speculated that the detection sensitivity of our 200x bulk sequencing is between 0.2% and 1%, and decided to have more unique L1 spiked at these two ratios. As a result, we spiked NA19240 at 0.2% and both HG00733 and HG00514 at 1%. (iii) NA12878 was chosen as the backbone in the mixing because it is from a homogeneous cell culture and is one of the most well-studied genomes.

The unique heterozygous MEIs in each of the spiked DNA samples are defined as: ${Unique\_MEI}_{i}={MEI}_{i}-\bigcup_{j=1,j\neq i}^{7} {MEI}_{j}$, where $i$ is one of the six DNA spiked at a ratio from 0.04% to 25%, and $j$ is one of six spiked DNA or NA12878 ($j=7$). For both of the mixed DNA (named “Mix”) and pure NA12878 (named “Control”), we made six separate libraries (TruSeq Nano) and sequenced each library to an average depth of 30-40x (total=200x). We applied RetroSom to call somatic MEIs that were found in the mixed DNA but not in the NA12878 control. The false positives and true positives were then defined as:

$${MEI}_{false\_positive}={MEI}_{Mix}-{MEI}_{Control}-\bigcup_{i=1}^{6} {MEI}_{i}$$

$${MEI}_{true\_positive\_i}={(MEI}_{Mix}-{MEI}_{Control})\bigcap{Unique\_MEI}_{i},$$

${MEI}_{Mix}$ is the set of MEIs called from the 200x sequencing of mixed DNA, ${MEI}_{Control}$ is the set of MEIs called from the 200x sequencing of NA12878 control, and $i$ is one of the six DNA spiked from 0.04% to 25%.

To evaluate the performance at different read depths, we down-sampled the sequencing data (“Mix” and “Control”) to 50x and 100x using Picard (DownsampleSam v2.17.3). We also mixed raw reads from previous sequencing data of each component at the same frequencies to create an *in silico*-mixing dataset of 200x, and combined it with the “Mix” sequencing data to a final depth of 400x. The sources include our own sequencing (A1S), HGSV dataset (HG00733, HG00514 and NA19238), BSMN common brain data, and the 200x Platinum Genomes dataset (NA12877). The 400x control data were created from combining the 200x NA12878 WGS in the Platinum Genomes and the “Control” sequencing data. Notably, we did not reuse the training data for testing at 400x depth, because RetroSom was initially trained on the 50x sequencing data of the 11 offspring (dbGAP: phs001224), not including the 200x sequencing data of their parents: NA12877 and NA12878 (ENA: PRJEB3246).

Postprocessing of putative somatic MEIs

RetroVis package to visualize the supporting reads

RetroSom includes a visualization tool, *RetroVis*, that systematically visualizes the supporting reads for each putative MEI with clear annotations for the insertion position, orientation, and other vital information (Extended Data Fig. 5A). Traditional genome browsers have issues with displaying the positions of both the anchor ends in the flanking sequences and the ME ends in the L1/*Alu* consensus. In addition, supporting reads for somatic MEIs are few in number and usually overwhelmed by other sequencing reads nearby.

In *RetroVis*, we annotate the human reference genome around the insertion junction as a black line on the top and the ME consensus on the bottom. The segment coordinates are labeled above the lines, and a short vertical line marks every 200 bases. Between them are the PE and SR supporting reads. Each PE supporting read is represented by a pair of arrows: a blue arrow and a red (or purple) arrow connected by a dashed line. The blue arrow represents the read that maps to flanking human genome sequences, and its location is based on the human reference on the top. The red (or purple) arrow represents the read that maps to the ME consensus, and its location is based on the ME consensus on the bottom. A red arrow indicates the MEI is inserted in the forward strand, while a purple arrow indicates the insertion is in the reverse strand. For the SR supporting read, the chimeric read that covers the insertion junction is plotted as a blue arrow connected to an empty rectangle. The blue arrow represents the read segment that maps to the flanking sequences, while the empty rectangle represents the ME segment, the alignment of which is indicated by a red/purple arrow below. This visualization provides a very convenient way to manually check any MEIs, especially when picking candidates for experimental validation.

Manual curation to remove false MEIs

To select a set of MEIs for experimental validation, we adopted a series of manual inspections to further eliminate likely false positives (Extended Data Fig. 5). We first examined the neighboring region of each putative MEI, removing novel junctions likely caused by structural variation, and regions with poor mapping quality (using the integrated genome browser, IGV)^15^. We also removed somatic MEIs present in datasets from other donors, likely occurring in regions prone to sequencing and mapping artifacts. We then used the visualization tool RetroVis to plot each insertion and its supporting reads, allowing for a rapid screening of multiple candidate MEI calls. Finally, we compared the sequences of the supporting reads to remove false insertions characterized by unexpected transduction, conflicting positions between support, or low homology in the ME ends mapped to the same location. The majority of the putative somatic MEIs were filtered during the manual curation, and the exact filters we used are listed in Supplementary Table 4.

Experimental validation of somatic MEIs

Probe-based droplet digital PCR (ddPCR)

All ddPCR assays were prepared using a published protocol^16^. The primer and probe sequences were designed with Primer3 (v2.3.7) on target templates from the SR supporting read or from stitching together pairs of PE supporting reads (Extended Data Figs. 6A and 7A). The primers and FAM-coupled ZEN double-quenched probes were synthesized at Integrated DNA Technologies (IDT). We used primers and a HEX-coupled probe for *RPP30* as the internal loading control, NA12878 genomic DNA as the negative control, and synthesized DNA oligo containing the insertion junction of interest as the positive control (IDT gBlocks gene fragments). Each candidate somatic MEI was analyzed in at least 4 replicates of genomic DNA from neurons, glia, and non-brain controls. Each replicate was incubated in a 20 μl reaction containing 30 ng genomic DNA, 0.9 μM primers for MEI junction, 0.9 μM primers for *RPP30*, 0.25 μM FAM probe (MEI junction), 0.25 μM HEX probe (*RPP30*), and 10 μl ddPCR supermix for probes (no dUTP). Sequences for the primers, probes, and gBlock controls are listed in Supplementary Table 5.

The reactions for L1 insertions were incubated as follows:

95°C for 10 min

94°C for 30 sec	|

59°C for 1 min	| 50 cycles

98°C for 10 min

The cutoffs separating the positive and negative droplets were chosen based on the negative and positive controls, and the levels of mosaicism were quantitated using QuantaSoft Analysis Pro Software(v1.0, BioRad). The target allele frequency is calculated from the number of positive droplets, based on the method described in Zhou et al. (2018)^16^. Under the assumption that somatic MEIs are heterozygous, their levels of mosaicism were calculated to be twice the allele frequency.

Nested PCR

We used two rounds of PCR to sequence the upstream and downstream junctions of the somatic MEIs (Extended Data Fig. 6A and Extended Data Fig. 7A). In the first PCR, we used primers on the flanking sequences surrounding the MEI and 60 ng genomic DNA. The pre-insertion allele is present in >99% of the cells and produces a strong band consistent with the coordinates in the human reference genome. The MEI-containing allele is expected to produce a larger product but is usually invisible on gel electrophoresis because of the amplification bias towards shorter and higher-frequency products. Nevertheless, we purified the DNA above the visible band from the first PCR, from a region that is 270-870 bp above for L1#1, or 260-610 bp­­­­­­­­­­­­­­­­­­­ above for L1#2 (Zymoclean Gel DNA recovery kit, Zymo research #D4007). In the second round of PCR, one half of the purified DNA was used to amplify the upstream junction using a primer in the upstream flanking sequence and a primer in the ME sequence. The other half was used to amplify the downstream junction using a primer in the downstream flanking sequence and a primer in the ME sequence. The nested PCR produced clean bands of expected size covering the upstream and downstream junctions, which were then analyzed with Sanger sequencing (Sequetech). Combining the junction sequences, we analyzed the exact MEI junction, target site duplications (TSD), endonuclease cutting sites, inserted ME sequences, and the microhomology between the ME sequence and the target site sequence if the L1 insertion was 5’-truncated. If there was a homology between the ME poly(A) tail and the TSD, we arbitrarily included the homologous region as part of the TSD (Extended Data Fig. 6D and Extended Data Fig. 7D)^17^.

All PCR reactions were incubated in a volume of 40 μl, containing 20 μl Phusion green Hotstart II HF PCR master mix (2x, Thermo Fisher), 0.9 μM of the primers, and the relevant template DNA. The primer sequences are in Supplementary Table 5. The reactions were incubated as follows:

95°C for 2 min

94°C for 45 sec	|

57°C (for L1#1) or 59°C (for L1#2) for 30 sec	| 30 cycles

72°C for 2 min	|

72°C for 7 min

Spatial distribution of L1#1 and L1#2

We sampled 11 additional pairs of tissues from symmetric regions in both hemispheres from the brain of donor 12004, including the (1) nearby STG, (2) superior frontal gyrus (marked as prefrontal cortex distal to STG), (3) inferior frontal gyrus (marked as prefrontal cortex proximal to STG), (4) motor cortex distal to STG, (5) motor cortex proximal to STG, (6) superior parietal lobule (marked as parietal cortex distal, (7) inferior parietal lobule (marked as parietal cortex proximal), (8) occipital cortex distal to STG, (9) occipital cortex proximal to STG, (10) putamen, and (11) cerebellum (Extended Data Fig. 9A). We separated the neurons and glial nuclei with FANS and used ddPCR to test for the presence and mosaicism of L1#1 and L1#2 in the genomic DNA of neurons and glia, respectively. Each DNA was tested in 4 technical replicate experiments using 30 ng of genomic DNA. The levels of mosaicism were calculated as twice the allele frequency, and we set the ddPCR detection threshold at >0.05% mosaicism (> 1 positive L1 junction droplet per replicate). The correlation between the mosaicism levels in neurons and in glia is shown in Extended Data Fig. 9F.

Reporter assay for L1#1 and L1#2

Extracting the full L1#1 and L1#2 sequences

Standard PCR across somatic L1 insertions fails to amplify the full L1 insertion sequences due to their lower levels of mosaicism and longer length compared to the pre-insertion alleles. Instead, we used overlap-extension PCR to stitch together the upstream and downstream junctions obtained from the nested PCR with a 17bp-overlap in the internal primers (Extended Data Fig. 9B)^18^. We first amplified 60 ng genomic DNA (12004 neuron) in two PCR reactions using external primers in the flanking sequences (primers **i** and **ii)**. We then cut out the blank gel region that was 270-870 bp above the pre-insertion allele product for L1#1 and 260-610 bp above the pre-insertion allele product for L1#2, and extracted DNA using the Zymoclean Gel DNA recovery kit (Zymo research #D4007). The blank gel contained the PCR product from the templates carrying the L1 insertions, and we eluted the extracted DNA in 13 μl water for each PCR reaction. We used 12.8 μl purified product in each nested PCR that amplified either the upstream (primers **iii** and **iv**) or downstream junctions (primers **v** and **vi**). For L1#1, a *Bam*HI site was attached to primer **iii**, and an *Apa*I site was attached to primer **vi**. Notably, because gene *FRMD4A* is in the reverse strand of the reference genome sequence, we attached a *Bam*HI site to primer **vi** and an *Apa*I site to primer **iii** for L1#2. There was a 17bp-overlap in the internal primers **iv** and **v**. We gel purified the upstream and downstream junctions using the Zymoclean Gel DNA recovery kit (Zymo research) and eluted the purified DNA in 10 μl water. The DNA concentration was quantified with Qubit (LifeTech cat# Q33216). Finally, we stitched together the two junctions in an overlap-extension PCR, using primers **iii** and **vi** and 100 ng of each junction. As a control for the genomic sequences without the L1 insertions, we amplified 60 ng NA12878 gDNA using primers **iii** and **vi** and purified the pre-insertion allele product from the introns of *CNNM2* and *FRMD4A*.

All PCR reactions were incubated in a volume of 40 μl, containing 20 μl Phusion green Hotstart II HF PCR master mix (2x, Thermo Fisher), 0.9 μM of the primer, and the relevant template DNA (60 ng for external PCR, 12.8 μl purified DNA for nested PCR, and 100ng of each junction for overlap-extension PCR). The primer sequences are in Supplementary Table 5. The reactions were incubated as follows:

95°C for 2 min

94°C for 45 sec	|

57°C (for L1#1) or 59°C (for L1#2) for 30 sec	| 30 cycles

72°C for 2 min	|

72°C for 7 min

Cloning into plasmid pGint

The L1#1 and L1#2, as well as the two control DNA, were digested using *Bam*HI-HF and *Apa*I enzymes (NEB# R3136S and R0114S, respectively). We first incubated 1 μg purified DNA with 1 μl *Apa*I enzyme and 3 μl NEB cutsmart (10X) buffer in a total volume of 29 μl for 2 hours at 25 °C. We then added 1 μl *Bam*HI-HF enzyme and incubated at 37 °C overnight. The reaction was stopped by adding 6 μl purple loading dye. The digested DNA was gel purified (Zymoclean Gel DNA recovery kit #D4007) and eluted in 10 μl water. The DNA concentration was quantified with Qubit (LifeTech cat# Q33216).

The DNA were ligated to the pGint plasmid using the instant sticky end ligase (2X) master mix (NEB M0370S) at 3-fold (insert DNA : vector) molar excess^19^. Specifically, *CNNM2* control (for L1#1) was mixed in a 15.5 μl reaction containing 37.5 ng control DNA, 85.5 ng pGint plasmid DNA, and 7.75 μl master mix. L1#1 was mixed in a 11.7 μl reaction containing 61.25 ng L1#1 DNA, 85.5 ng pGint plasmid DNA, and 5.85 μl master mix. *FRMD4A* control (for L1#2) was mixed in a 10 μl reaction containing 12.5 ng control DNA, 80 ng pGint plasmid DNA, and 5 μl master mix. L1#2 was mixed in a 10 μl reaction containing 36.3 ng L1#2 DNA, 80 ng pGint plasmid DNA, and 5 μl master mix. At the same time, we also prepared the vector-only controls using only 85.5 ng (for L1#1 and control) or 80 ng (for L1#2 and control) pGint plasmid DNA. The ligation reaction was mixed and left on ice for 5 min.

For each cloning experiment, we thawed 50 μl TOP10 competent cells on ice and incubated them on ice for 30 min with a 2 μl ligation reaction. We then heat shocked the cells at 42 °C for 30 sec, put them back on ice for 2 min, and recovered them into 950 μl SOC media (Invitrogen #15544-034) at 37 °C for 1 hour. We plated the cells on Kan-50 selection plates (Teknova) overnight at 37 °C.

We verified whether the colonies contained the correct insert with colony PCR and Sanger sequencing. We picked single colonies from the Kan-50 selection plates and spiked each colony in 5ml LB (Teknova L8000) culture with 25 μl Kan50 (10mg/ml). In the colony PCR, we tested each colony in a 20 μl PCR reaction containing 1 μl of the LB culture, 10 μl Phusion green Hotstart II HF PCR master mix (2x, Thermo Fisher), and 0.9 μM of primer (Supplementary Table 5). The reactions were incubated as follows:

95°C for 2 min

94°C for 45 sec	|

55°C for 30 sec	| 30 cycles

72°C for 2 min	|

72°C for 4 min

We examined the PCR product on gel electrophoresis to check for the correct insert size. In addition, we incubated the 5ml LB culture at 37 °C overnight with shaking. We extracted the plasmid using miniprep (Qiagen Cat #27106) and verified the insert sequence with Sanger sequencing. The validated clones were named as follows: GL1#1 (1123bp insert with L1#1 and flank), GL1#2 (686bp insert with L1#2 and flank), Gcont#1 (691bp insert with flanking sequence for L1#1), and Gcont#2 (240bp insert with flanking sequence for L1#2).

Transient transfection of reporter plasmids into HeLa cells

The four plasmids, Gcont#1, GL1#1, Gcont#2, and GL1#2, were transfected into HeLa S3 cells with Lipofectamine 3000 reagent in two separate experiments: (i) dual transfection together with a red fluorescence protein reporter (RFP) ‘Rint’ in 5 wells per reporter and (ii) single transfection without Rint in 2 wells per reporter. For the dual-transfection experiment, we started by seeding HeLa cells on a 24-well plate (~70% confluence, 50,000 cells per well). On the next day, we prepared the Lipofectamine mixture containing 33.75 μl Lipofectamine 3000 (Thermo Fisher) and 562.5 μl Opti-MEM media (Thermo Fisher). We then prepared a plasmid DNA mixture for each reporter: (i) 4.36 μl Gcont#1 plasmid (375 ng/μl) with 1.95 μl Rint plasmid (900 ng/μl), 145.4 μl Opti-MEM media, and 6 μl P3000 reagent; (ii) 6.64 μl GL1#1 plasmid (266 ng/μl) with 1.95 μl Rint plasmid, 147.6 μl Opti-MEM media, and 6 μl P3000 reagent; (iii) 1.01 μl Gcont#2 plasmid (1480 ng/μl) with 1.95 μl Rint plasmid, 150 μl Opti-MEM media, and 6 μl P3000 reagent; (iv) 1.10 μl GL1#2 plasmid (1480 ng/μl) with 1.95 μl Rint plasmid, 150 μl Opti-MEM media, and 6 μl P3000 reagent. The same number of copies of plasmids was used in each mixture, as the amount was calculated based on the plasmid size: Gcont#1=5410 bp, GL1#1=5482 bp, Gcont#2=4959 bp, GL1#2=5405 bp, and Rint=5816 bp. For each plasmid, we mixed 133.75 μl Lipofectamine mixture with 133.75 μl plasmid mixture, incubated the mixture at room temperature for 15 min, and applied 50 μl to each of the 5 wells. The order of each transporter assay was shuffled and kept hidden until the fluorescence was quantitated by a different experimenter to allow for a blind experiment.

Similar protocol was used in the single-transfection experiment, except for the plasmid mixtures (prepared for 2.25 reactions): (i) 0.787 μl Gcont#1 plasmid (780 ng/μl) with 56.25 μl Opti-MEM media and 1.125 μl P3000 reagent; (ii) 0.745 μl GL1#1 plasmid (890 ng/μl) with 56.25 μl Opti-MEM media and 1.125 μl P3000 reagent; (iii) 0.380 μl Gcont#2 plasmid (1480 ng/μl) with 56.25 μl Opti-MEM media and 1.125 μl P3000 reagent; (iv) 0.416 μl GL1#2 plasmid (1480 ng/μl) with 56.25 μl Opti-MEM media and 1.125 μl P3000 reagent.

Fluorescence quantification

After incubating HeLa cells with the transfection mixtures for 23 hours, we captured images in GFP, RFP, and bright field channels (Leica DMI 3000B) in each well on the top-center, bottom-left, and bottom-right sections (Fig. 4E, F and Extended Data Fig. 9C). The GFP and RFP images were taken with an exposure time of 200ms and analog gain of 9, and the bright field images were taken with an exposure time of 40ms and analog gain of 2. We also confirmed, in a separate pilot experiment, the absence of bleed-through interference between the GFP and RFP channels.

On each image, we labeled all cells with visible fluorescence signals (green or red) with a region of interest (ROI) marker that were adjusted to fit the cell shape, as well as five blank regions (top-left, top-right, center, bottom-left, bottom-right), to measure the background fluorescence (Leica Application Suite 300 build 8134). We used $\overline{ROI}-\overline{Background}$ to represent the signal strength of each cell, where $\overline{ROI}$ represents the mean intensity value of pixels in ROI, and $\overline{Background}$ represents the mean intensity value in all five blank regions. We excluded dead/broken cells and image artifacts by referring to the bright-field image. The number of plasmids transfected into each cell is highly variable but the impact from each reporter can be evaluated after averaging a large number of cells. From 2 independent experiments and 7 wells per plasmid, we quantitated a total of 912 cells for GCont#1, 785 cells for GL1#1, 878 cells for GCont#1, and 701 cells for GL1#2 before the plasmid labels were revealed for statistical analysis.

Estimation of poly(A) tail sizes

We evaluated the length of poly(A) tails in 24 GL1#1 clones and 24 GL1#2 clones using Sanger sequencing (Extended Data Fig. 9E). The variable poly(A) tails are likely caused by polymerase slippage around low complexity sequences, leading to both longer and shorter poly(A) sizes^20^. We chose the size supported by the highest number of clones as the estimates for the poly(A) length. Our estimations of poly(A) sizes required PCR amplification from the tissue DNA and may have introduced biases towards shorter products and templates with higher mosaicism^21^.

Statistical analysis

We used Student’s *t*-test (unpaired, equal variance) to calculate the statistical significance of the mosaicism difference in various fractions (Extended Data Figs. 6C and 7C). The correlation of L1 mosaicism levels between neurons and glia in different anatomical regions is evaluated by rank-based Spearman ρ statistic (Extended Data Fig. 9F).

To evaluate the level of fluorescence in the transfection experiments (Fig. 4C, D and Extended Data Fig. 9D), we performed a log transformation on the fluorescence intensities and then used Student’s *t*-test to compare the overall levels of fluorescence between groups. A dummy variable was added to all fluorescence values to remove 0 and negative values, and the log transformation was to make the intensity values closer to a normal distribution. We performed 10 statistical tests comparing various groups in the transfection experiments and adjusted the p value using the Bonferroni correction: $adjusted\_p\_value=10*p\_value$.

A possible explanation for the lower fluorescence level in L1 reporters compared to that in controls is slower transcription due to larger $insert\_length$ (Fig. 4C and Extended Data Fig. 9D). However, our data suggest that the difference in the tested range of $insert\_length$ (240 to 1123bp) is unlikely to be the only contributing factor to the difference between L1 and control reporters. The fluorescence in Gcont#1 (686bp) is similar to that in Gcont#2 (240bp) (adjusted *p*=1) but significantly stronger than that in GL1#2 (691bp) (adjusted *p*=2.6x10^-22^). In addition, the fluorescence in GL1#1 is stronger than in GL1#2 (adjusted *p*=3x10^-4^), despite larger $insert\_length$ (1123bp vs. 691bp).

To further evaluate the impact of insert size, we built a linear regression model to fit the GFP fluorescence for all four plasmids: $log\left( GFP \right)\sim L1+log\left( insert\_length \right)$, where $L1$ is a binary variable indicating whether the plasmid has an L1 insertion $\left( L1=1 \right)$ or is a control $\left( L1=0 \right)$, and $insert\_length$ is 691 for Gcont#1, 1123 for GL1#1, 240 for Gcont#2, and 686 for GL1#2. In this linear model, the $insert\_length$ does not affect the fluorescence intensity significantly (adjusted *p*=0.25, coefficient=0.10), while $L1$ is negatively correlated with the fluorescence intensity (adjusted *p*=4x10^-19^, coefficient=-0.485).

EXTENDED DATA

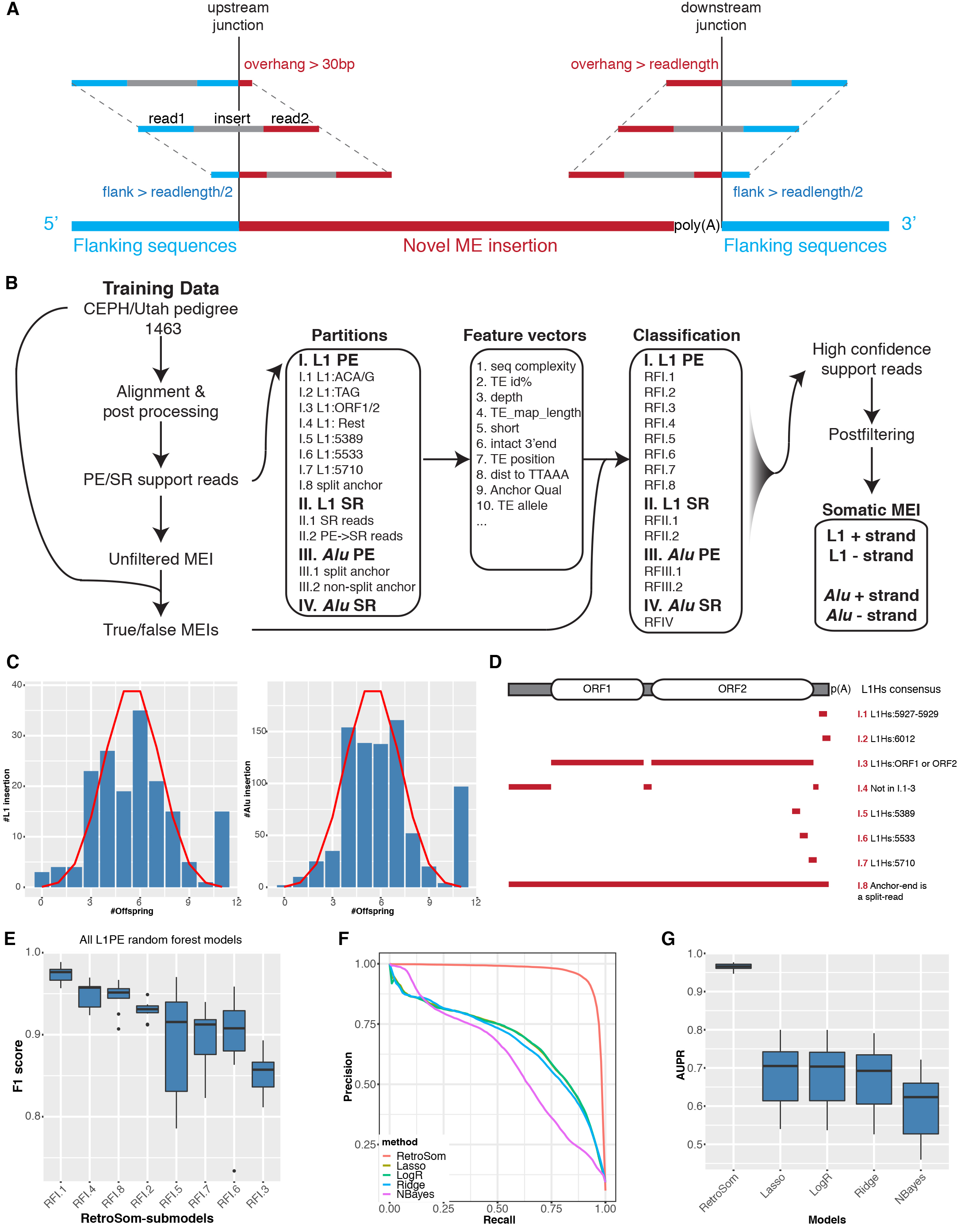

**Fig. 1. Classification of supporting reads from putative mobile element insertions.**

**A**) We simulated the relationship between the detectable mosaicism of somatic MEIs and the number of supporting reads in bulk sequencing by considering the range of coordinates for the putative supporting reads for either the upstream or downstream junction (see Fig. 1D). Blue, segment of supporting read that maps to flanking sequence; red, segment of read that maps to ME consensus; gray, the insert segment between the two paired-end reads.

(**B**) A detailed flowchart describing the framework behind RetroSom. We labeled putative supporting reads as true or false insertions based on the inheritance pattern and built a set of random forest models to classify them based on various sequencing features (see Supplementary Table 3).

(**C**) The distribution of true L1 (left) and *Alu* (right) insertions among 11 offspring is similar to a theoretical binomial distribution (red line). The peaks around N=11 represent additional MEIs that are homozygous in one of the parents and transmitted to all 11 offspring.

(**D**) To avoid missing values, we categorized L1 PE supporting reads into 8 subgroups depending on their mapping locations on the L1Hs (L1 human specific) consensus sequence.

(**E**) The performance of random forest classification in all 8 L1 PE read sub-models, ranked based on their average F1 score (harmonic average of sensitivity and precision).

(**F and G**) Model selection and evaluation with 11x cross validation: (**F**) precision-recall curve, (**G**) area under the precision-recall curve (AUPR).

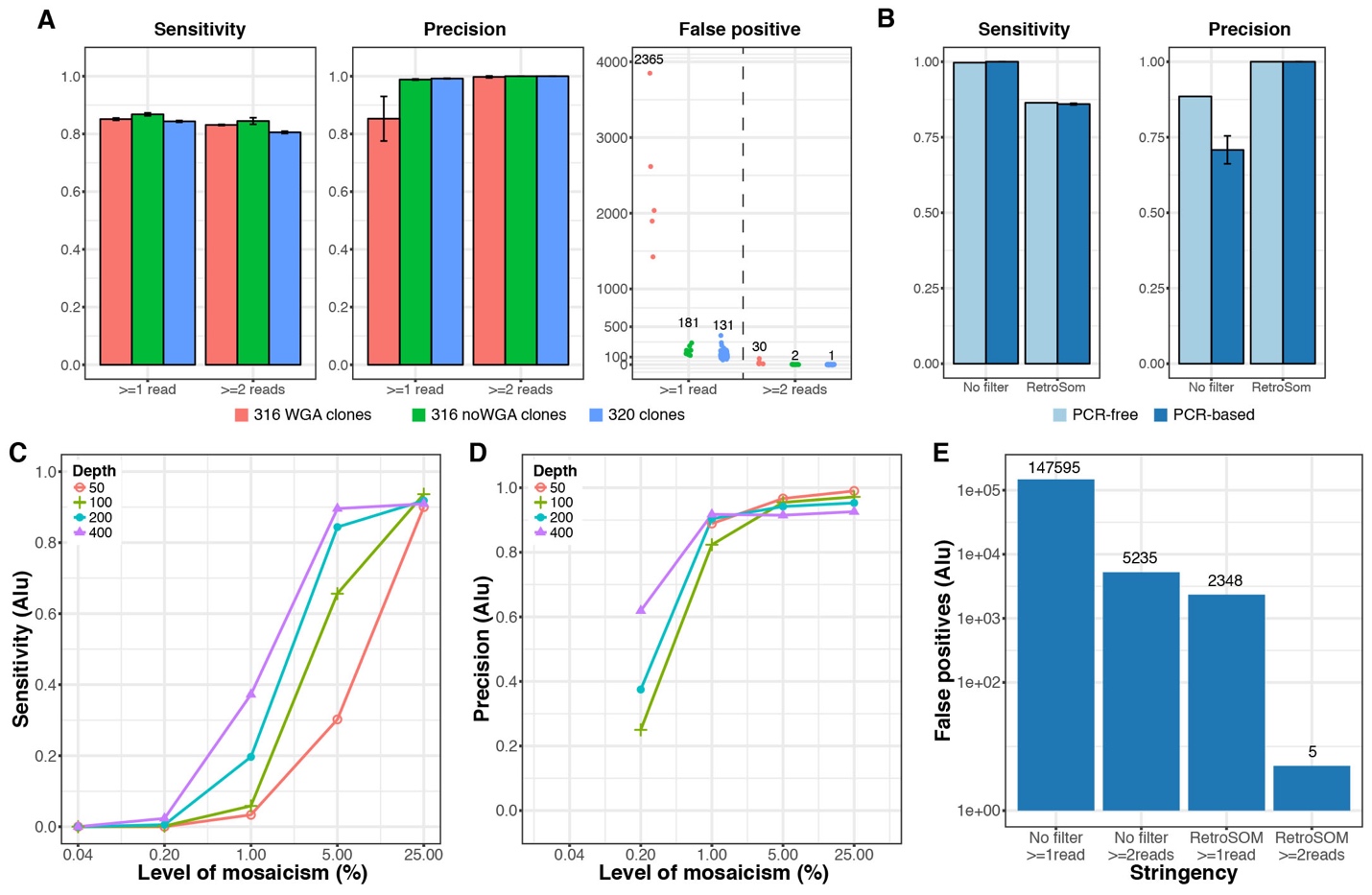

**Fig. 2:** **Benchmarking *Alu* insertions in independent test datasets.**

(**A**) Performance in detecting germline *Alu* insertions from clonally expanded fetal brain cells sequencing data. Red, clones from donor “316” sequenced with whole genome amplification (316WGA); green, the rest of the “316” datasets (316 noWGA); blue, clones from donor “320”. The error bars represent the 95% confidence intervals estimated in individual clones.

(**B**) Performance in detecting germline *Alu* insertions from sequencing libraries prepared with or without PCR. A similar comparison for libraries created with A1S heart tissue produced similar results (Extended Data Fig. 4).

(**C-E**) Performance in detecting somatic MEIs simulated by six genomic DNA samples at proportions of 0.04% to 25% with that of NA12878, at various sequencing depth (red, 50$\times$; green, 100$\times$; blue, 200$\times$; purple, 400$\times$).

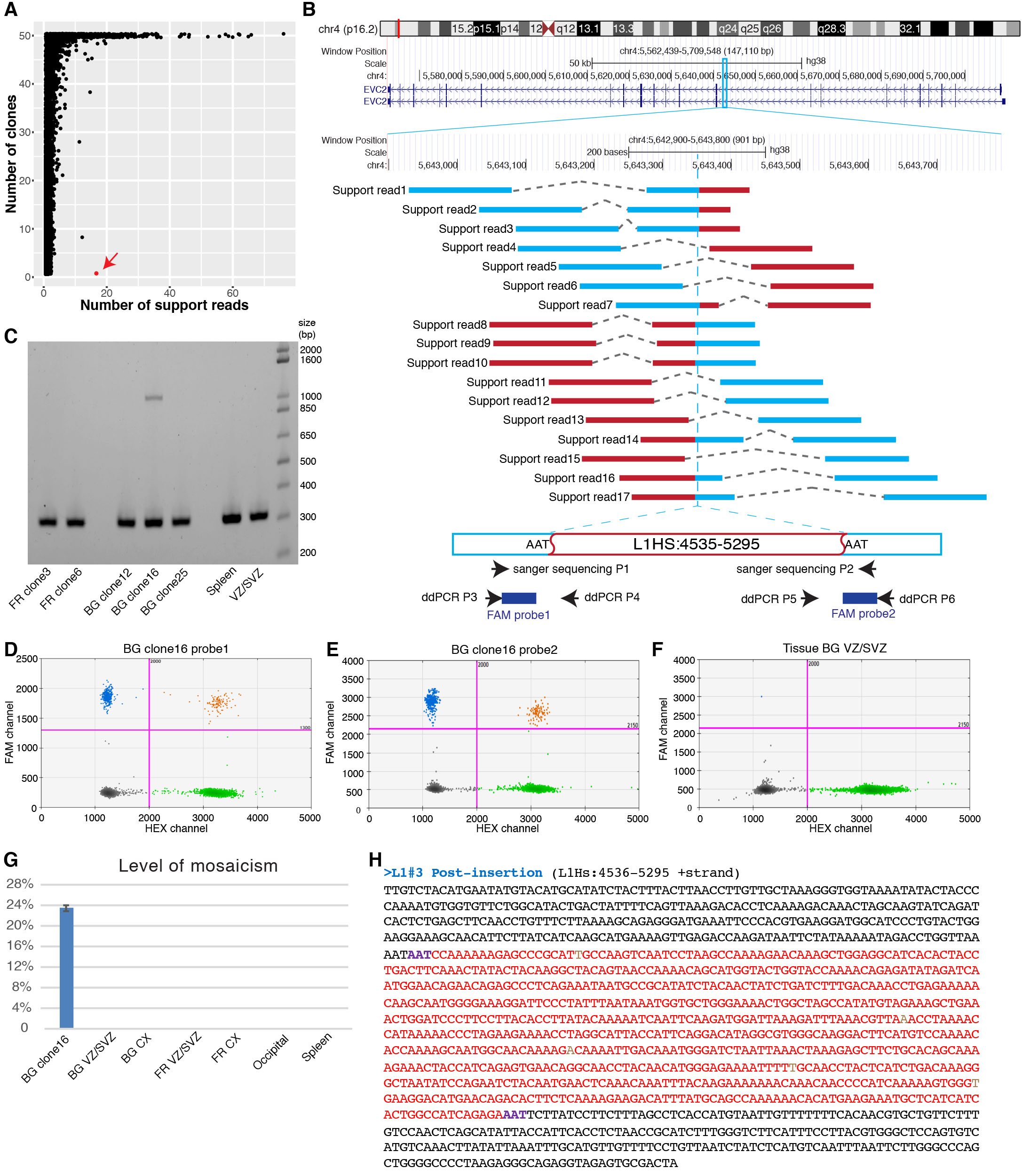

**Fig. 3­­­. Discovery and experimental validation of insertion L1#3.**

(**A**) We identified a somatic L1 insertion (L1#3, red arrow) in one clone, “BG clone16,” with 17 supporting reads.

(**B**) L1#3 is inserted into an intron of gene *EVC2*. Blue, segment of supporting read that maps to the flanking sequence; red, segment of read that maps to ME consensus.

(**C**) PCR surrounding L1#3 produced a unique band in BG clone16, as well as a lower band in all tested samples, representing the product from the DNA without the insertion.

(**D**) DdPCR detects the upstream junction in 22.54% of the cells in BG clone16.

(**E**) DdPCR detects the downstream junction in 24.16% of the cells in BG clone16.

(**F and G**) L1#3 is absent in 6 bulk tissues: BG ventricular zone/subventricular zone (BG VZ/SVZ), BG cortex (BG CX), FR VZ/SVZ, FR CX, occipital cortex, and spleen.

(**H**) The full sequence of L1#3: black, flanking sequence; red, inserted L1 sequence; purple, target site duplication; brown, mismatches to the L1Hs consensus.

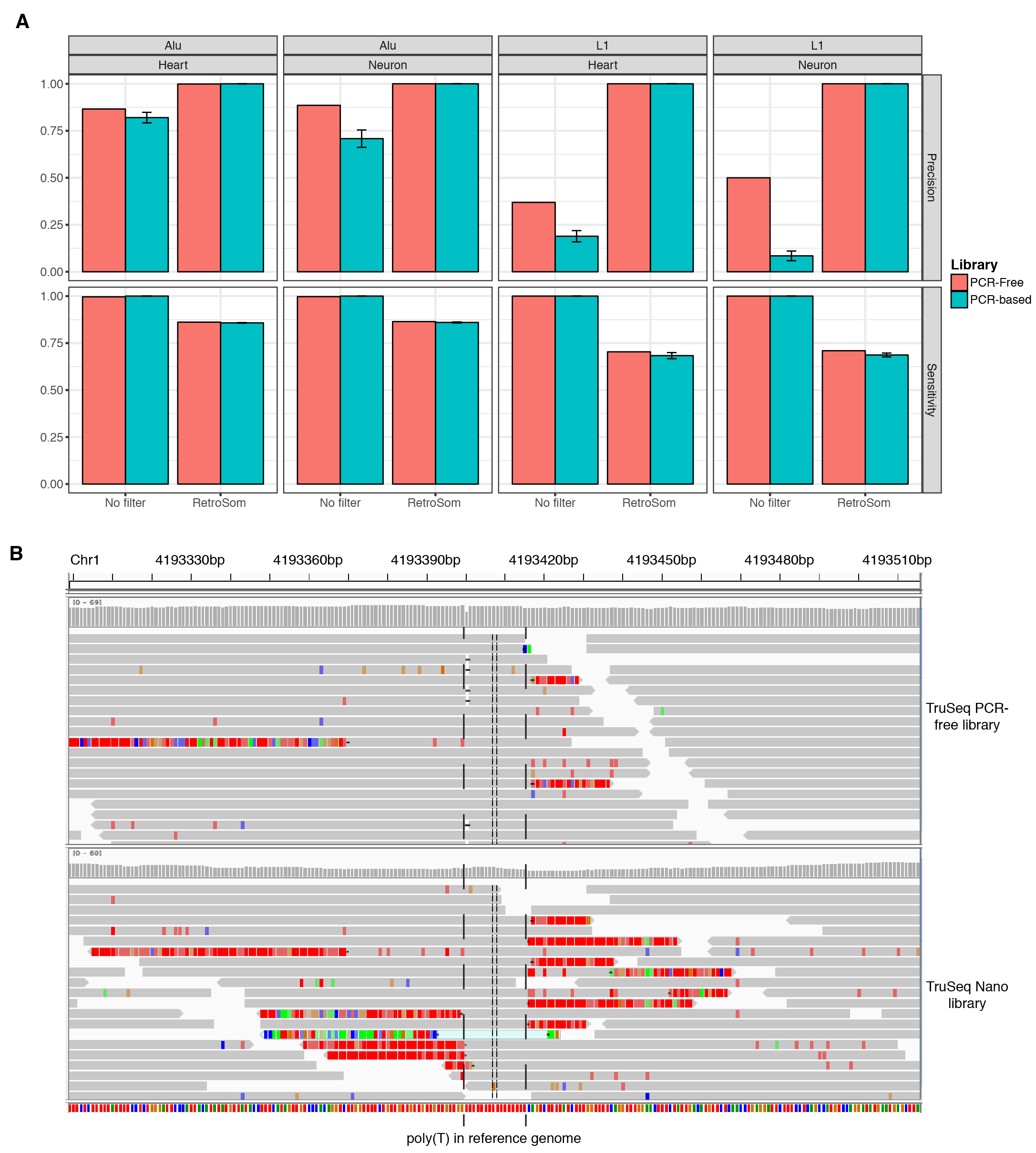

**Fig. 4. Comparison between Illumina TruSeq PCR-free libraries and PCR-based libraries.**

(**A**) Comparison of MEI detection precision (top row) and sensitivity (bottom row) for *Alu* (1st and 2nd columns) and L1 (3rd and 4th columns) between the PCR-free (red) and the PCR-based (green) sequencing libraries for A1S heart cells and neurons.

(**B**) Examples of sequencing errors introduced by PCR. At a locus with a 16bp poly(T) sequence, most of the reads from the PCR-free library are properly mapped (solid gray box), while the reads from the PCR-based TruSeq Nano library tend to have many sequencing errors, indicated by mismatches (green, A; blue, C; brown, G; red, T) and inserts of additional T.

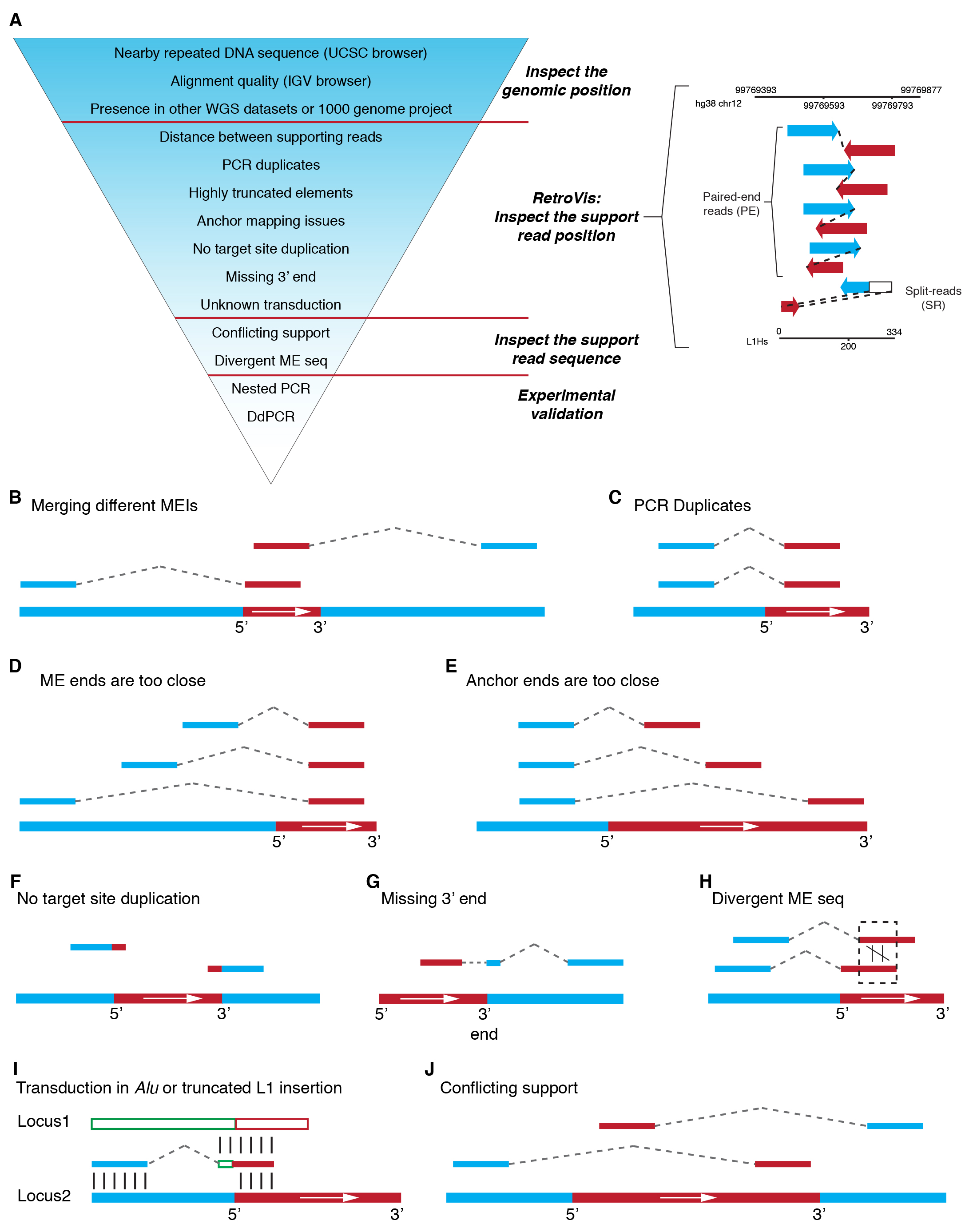

**Fig. 5. Postprocessing of putative somatic MEIs.**

(**A**) Procedure for manual curation of putative somatic MEIs. To further remove false positive MEIs, especially for *Alu* insertions, we implement manual inspections for each putative insertion. We first check the neighboring regions in both the UCSC and IGV browsers and remove calls that are from regions of potential mapping errors or CNVs. We also remove calls that are found in datasets of other donors. We then apply a novel visualization tool, *RetroVis*, to quickly screen out calls with questionable supporting read positions. We further inspect the read sequences to check for unwarranted transduction and similarity between different supporting reads. Finally, we design nested PCR and ddPCR to validate the insertions and quantify their respective levels of mosaicism using DNA from the same tissue.

In a *RetroVis* plot, black lines represent human genome location (top) and the inferred segment of the inserted mobile element (e.g., L1) (bottom). A paired-end supporting read is represented by a blue arrow and a red (+ strand insertion) or purple (-strand insertion) arrow connected by a dashed line. A split-read supporting read (spanning an insertion junction) is plotted as a blue arrow (reference segment) connected to an empty rectangle (mobile element segment), with a red or purple arrow below. The positions of the blue segments and red/purple segments reflect the insertion coordinates in the human reference genome and mobile element consensus.

(**B-J**) Examples of likely false positive insertions examined by manual curation. Blue, flanking sequence; red, mobile element sequence (+ strand insertion).

(**B**) Merging different MEIs into one.

(**C**) PCR duplicates.

(**D**) All ME ends are mapped to identical coordinates at the 3’ end of the L1Hs sequence.

(**E**) All anchor ends are mapped to identical coordinates in flanking sequences.

(**F**) Lacking target site duplication.

(**G**) A truncated 3’ end indicates a false insertion or an endonuclease-independent retrotransposition.

(**H**) Two supporting reads mapping to the same ME location but having a low sequence similarity.

(**I**) When the split-read supporting read is mapped partially to the ME consensus (red, locus 2) and fully to another reference genome element (green and red, locus 1), the additional sequence (green) is transduced to the new location. Transduction in *Alu* insertions, or 5’ transduction in 5’-truncated L1 insertions, indicates a false insertion.

(**J**) The supporting reads suggest that the ME is inserted in the + strand, yet the 3’ end is closer to the upstream flank and the 5’ end is closer to the downstream flank. This conflict indicates a false insertion or a 5’ inversion in L1 retrotransposition.

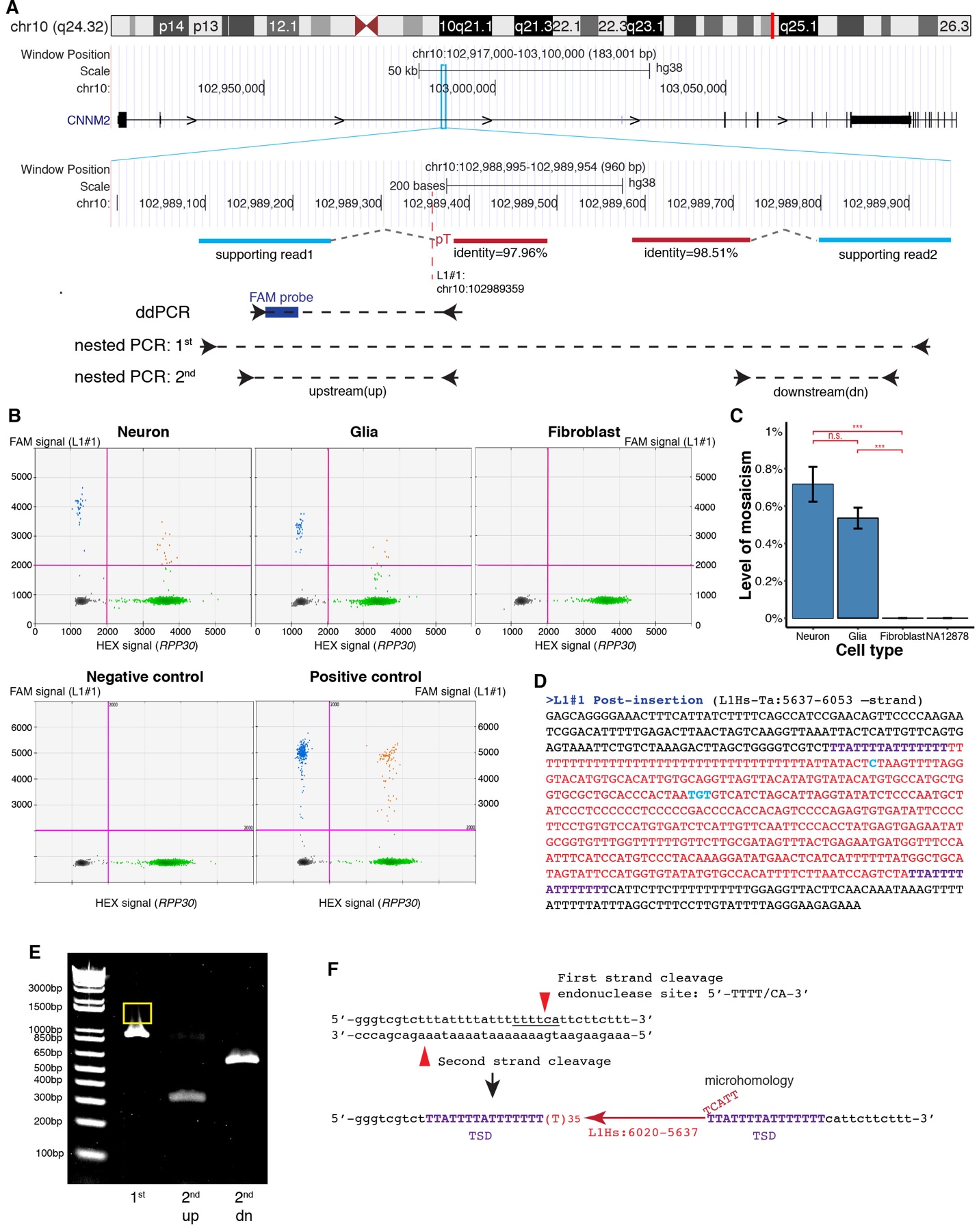

**Fig. 6. Validation of L1#1.**

(**A**) L1#1 was identified by RetroSom with two supporting sequencing reads, and the insertion is in the antisense strand of an intron of gene *CNNM2*^22^. Blue, read that maps to the flanking sequence; red, mate read that maps to the L1 consensus. We targeted putative L1 insertion junctions with ddPCR to quantitate their levels of mosaicism and used nested PCR to determine the junction sequences.

(**B**) DdPCR detected a clear signal for L1#1 in the brain, in both neurons and glia, but not in the fibroblast. Green, droplets containing only *RPP30* (internal control); Blue, droplets containing only the L1 junction template; Orange, droplets containing both L1 and *RPP30* templates; Black, droplets containing neither L1 nor *RPP30* templates. We used NA12878 DNA as a negative control and synthesized DNA with the target L1 junction as a positive control.

(**C**) DdPCR confirmed that L1#1 is present in both neurons (0.72%) and glia (0.54%), and absent in the fibroblast and NA12878. n.s.,**,*** indicates *p* value (Student’s t-test) > 0.05 or < 0.01, 0.001, respectively. The error bars represent the 95% confidence intervals.

(**D**) The full sequences of L1#1. Black, flanking sequence; red, inserted L1 sequence; purple, target site duplication; cyan, L1Hs specific alleles; brown, mismatch to the L1Hs consensus.

**(E**) Nested PCR for L1#1. We used the DNA product above the band produced by the pre-insertion DNA (yellow rectangle) as the template for the subsequent PCRs, which amplify the upstream (2^nd^ up) and downstream (2^nd^ dn) junctions of L1#1 specifically.

(**F**) L1#1 has an endonuclease cleavage site 5’-TTTT/CA-3’ and a 15bp TSD. The inserted L1 element is truncated on the 5’ end, with a microhomology between the L1 sequence and the target site.

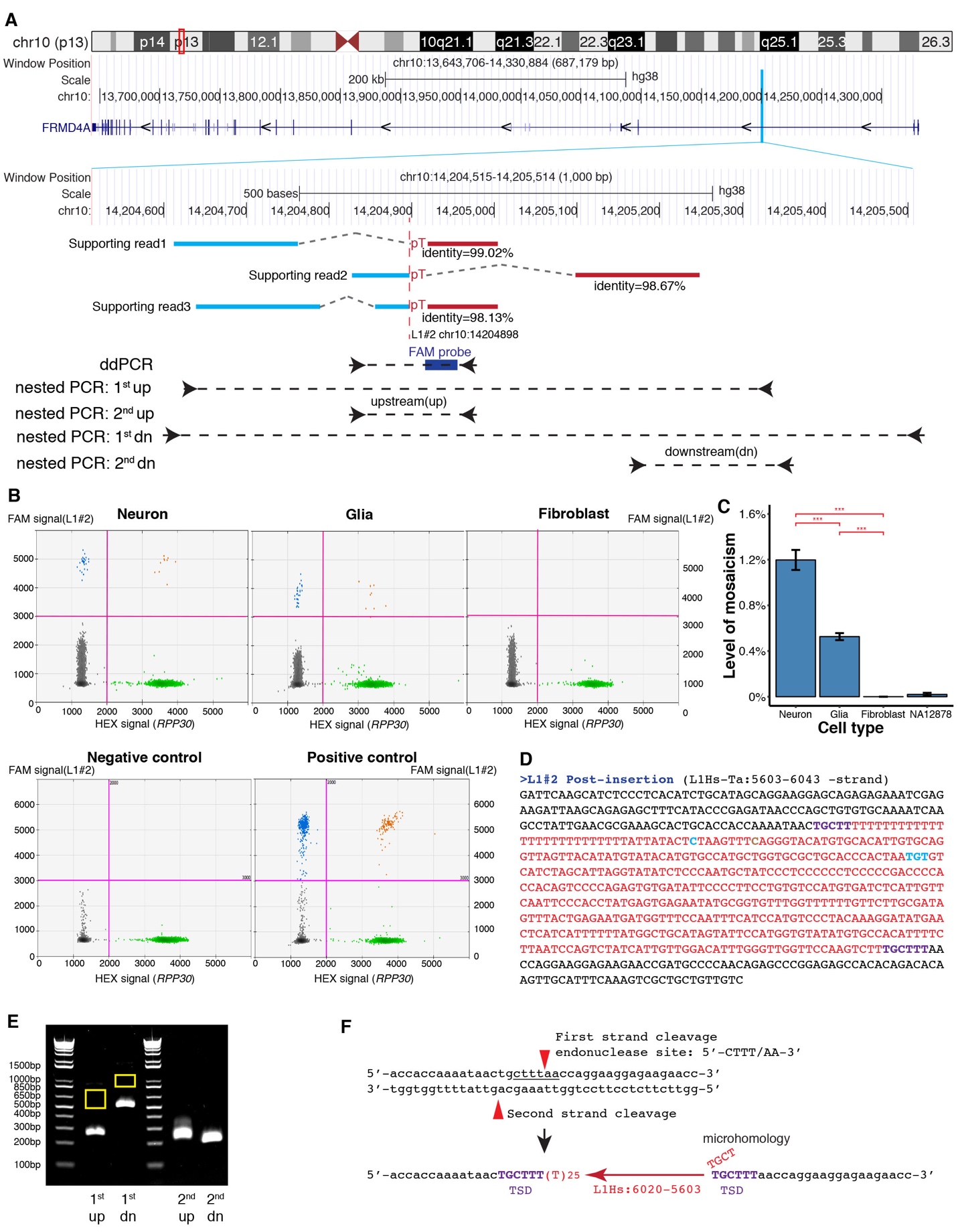

**Fig. 7. Validation of L1#2.**

(**A**) L1#2 was identified by RetroSom with three supporting sequencing reads, and the insertion is in the sense strand of an intron of gene *FRMD4A*. We targeted putative L1 insertion junctions with ddPCR to quantitate their levels of mosaicism and used nested PCR to determine the junction sequences.

(**B**) DdPCR detected a clear signal for L1#2 in the brain, in both neurons and glia, but not in the fibroblast. Green, droplets containing only *RPP30* (internal control); Blue, droplets containing only the L1 junction template; Orange, droplets containing both L1 and *RPP30* templates; Black, droplets containing neither L1 nor *RPP30* templates. We used NA12878 DNA as a negative control and synthesized DNA with the target L1 junction as a positive control.

(**C**) DdPCR confirmed L1#2 is present in both neurons (1.2%) and glia (0.53%), and absent in the fibroblast and NA12878. n.s.,**,*** indicates *p* value (Student’s t-test) > 0.05 or < 0.01, 0.001, respectively. The error bars represent the 95% confidence intervals.

(**D**) The full sequences of L1#2. Black, flanking sequence; red, inserted L1 sequence; purple, target site duplication; cyan, L1Hs specific alleles; brown, mismatch to the L1Hs consensus.

**(E**) Nested PCR for L1#2. We used the DNA product above the band produced by the pre-insertion DNA (yellow rectangles) as the template for the subsequent PCRs, which amplify the upstream (2^nd^ up) and downstream (2^nd^ dn) junctions of L1#2 specifically.

(**F**) L1#2 has an endonuclease cleavage site 5’-CTTT/AA-3’ and a 6bp TSD. The inserted L1 element is truncated on the 5’ end, with a microhomology between the L1 sequence and the target site.

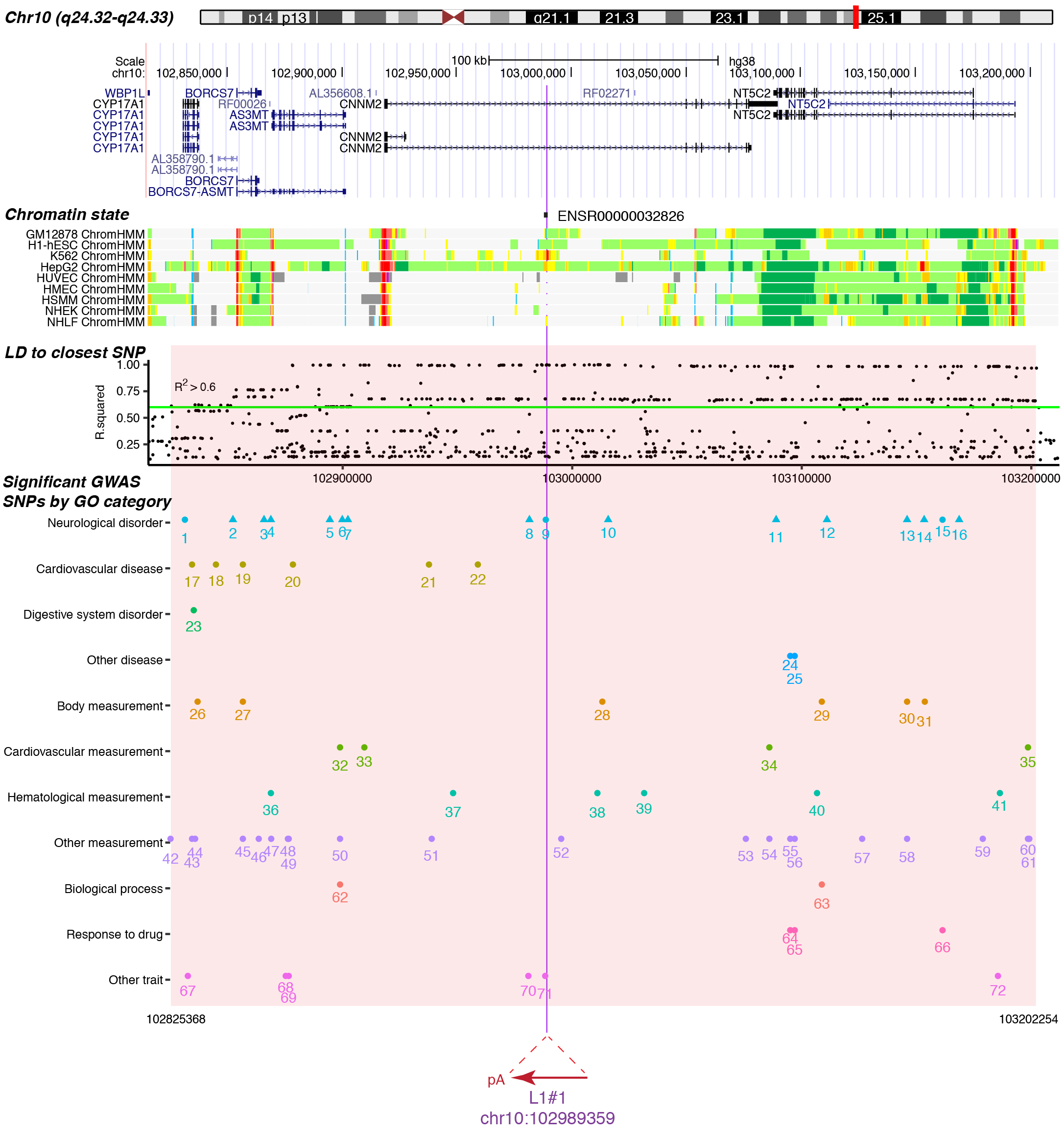

**Fig. 8. The genomic locus with L1#1 insertion.**

L1#1 is inserted in a 1639bp promoter flanking region (ENSR00000032826) that is hypothesized to regulates the expression of nearby genes^23^. The chromatin states are shown for a subset of human cell lines: light gray, heterochromatin; light green, weakly transcribed; yellow, weak/poised enhancer; orange, strong enhancer; light red, weak promoter; bright red, strong promoter. A comprehensive list of all evidence is in Supplementary Table 6. L1#1 is inserted in a linkage disequilibrium (LD) block, based on the common SNPs that are highly correlated (R2 > 0.6, green line) with the closest common SNP to L1#1, rs1890185. This LD block is highlighted in red, and contains 72 lead SNPs associated with 10 diseases or disorders and 28 measurements or other traits^24^, including 13 risk SNPs from 11 schizophrenia studies (triangle). We categorized all traits under 11 terms based on the Experimental Factor Ontology^25^. The significant SNPs, indexed from number 1 to 72, are documented in details in Supplementary Table 7.

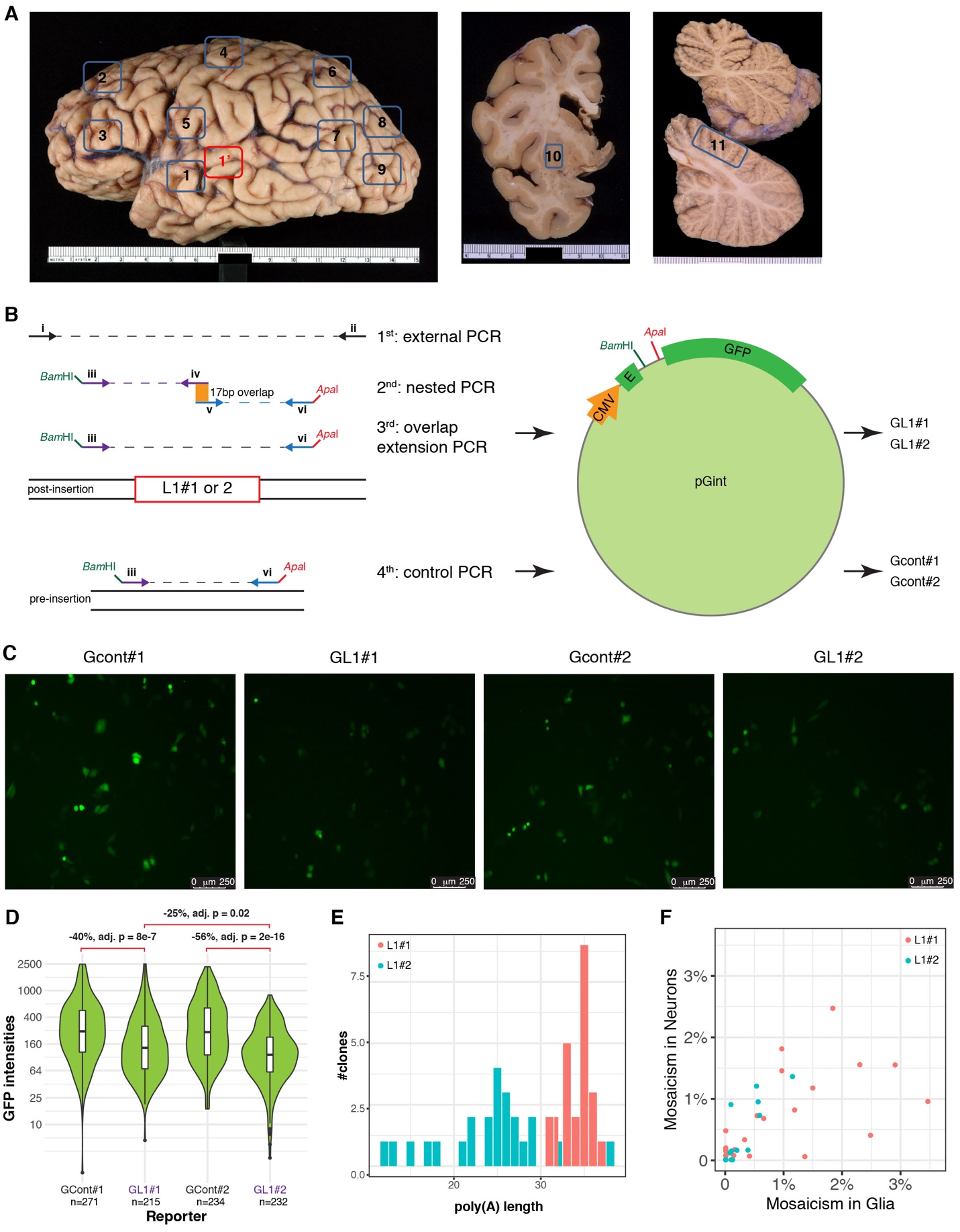

**Fig. 9. Spatial and functional analysis of somatic L1 insertions L1#1 and L1#2.**

(**A**) Anatomical brain regions studied in donor 12004: 1’, right superior temporal gyrus for whole genome sequencing; 1, superior temporal gyrus (both sides); 2, prefrontal cortex distal (superior frontal gyrus, both sides); 3, prefrontal cortex proximal (inferior frontal gyrus, both sides); 4, motor cortex distal (both sides); 5, motor cortex proximal (both sides); 6, parietal cortex distal (superior parietal lobule, both sides); 7, parietal cortex proximal (Inferior parietal lobule, both sides); 8, occipital cortex distal (both sides); 9, occipital cortex proximal (both sides); 10, putamen (both sides); 11, cerebellum (both sides). The tissues that were dissected from both hemispheres were bilaterally symmetrical. The metric unit on the ruler is the centimeter.

(**B**) Cloning the L1#1, L1#2, and controls to plasmid Gint. We used nested PCR to amplify the 5’ and 3’ junctions for L1#1 and L1#2 and overlap extension PCR to obtain the full sequence of L1#1 and L1#2. Control DNA was amplified on DNA without the L1 insertion (NA12878) using primer iii and primer vi. The amplified DNA (L1 or control) was cloned to a constitutively spliced intron in an enhanced green fluorescence protein (EGFP) reporter, pGint.

(**C**) GFP fluorescence of the control and L1#1 reporters in the single transfection experiment (no Rint).

(**D**) L1 reporters also reduced fluorescence significantly in experiment (**C)**, with a stronger effect in L1#2 than in L1#1.

(**E**) Poly(A) lengths of L1#1 and L1#2 were estimated as the lengths supported by the highest numbers of GL1#1 and GL1#2 clones. Red, L1#1; green, L1#2. The variation among clones was likely the result of PCR stutter around low-complexity templates^20^.

(**F**) The levels of mosaicism in neurons are highly correlated with levels in glia. Red, L1#1; green, L1#2.

Supplementary Tables

**Supplementary Table 1. List of studied specimens.**

| **Donor**  **ID** | **A1S** | **10011** | **11003** | **11004** | **12004** | **F1** |
| --- | --- | --- | --- | --- | --- | --- |
| **CODE** | A34605 | CT-CF-10-011 | CT-CF-11-003 | CT-CF-11-004 | CT-CF-12-004 | F7030320151530 |
| **Age** | 80 y.o. | 47 y.o. | 55 y.o. | 49 y.o. | 45 y.o. | 18.3 weeks |
| **Sex** | Male | Male | Male | Male | Male | unknown |
| **Race** | Caucasian | Caucasian | Caucasian | Caucasian | Caucasian | African American |
| **PMI** | 9 | 23 | 24 | 24 | 25.6 | unknown |
| **Matched**  **Pair** | N.A. | **12004** | **11004** | **11003** | **10011** | N.A. |
| **Tissue** | NeuN+, NeuN-, Heart | NeuN+, NeuN-, Fibroblast | NeuN+, NeuN-, Fibroblast | NeuN+, NeuN-, Fibroblast | NeuN+, NeuN-, Fibroblast | Neurons, Astrocytes, Heart |
| **DIN** | >7.0 | >7.0 | >7.0 | >7.0 | >7.0 | >7.0 |
| **SCZ** | No | No | No | Yes | Yes | N.A. |

**Supplementary Table 2. List of whole genome sequencing datasets.**

| **ID** | **Tissue** | **Source** | **Depth (X)** | **Library Type** | **Read length** | **Purpose** |
| --- | --- | --- | --- | --- | --- | --- |
| A1S_Heart | Heart | this project | 200 | TruSeq Nano | 2x150bp | Discovery |
| A1S_Neuron | STG NeuN+ | this project | 200 | TruSeq Nano | 2x150bp | Discovery |
| A1S_Glia | STG NeuN- | this project | 200 | TruSeq Nano | 2x150bp | Discovery |
| 10011_Fib | Fibroblast | this project | 200 | TruSeq Nano | 2x150bp | Discovery |
| 10011_Neuron | STG NeuN+ | this project | 200 | TruSeq Nano | 2x150bp | Discovery |
| 10011_Glia | STG NeuN- | this project | 200 | TruSeq Nano | 2x150bp | Discovery |
| 11003_Fib | Fibroblast | this project | 200 | TruSeq Nano | 2x150bp | Discovery |
| 11003_Neuron | STG NeuN+ | this project | 200 | TruSeq Nano | 2x150bp | Discovery |
| 11003_Glia | STG NeuN- | this project | 200 | TruSeq Nano | 2x150bp | Discovery |
| 11004_Fib | Fibroblast | this project | 200 | TruSeq Nano | 2x150bp | Discovery |
| 11004_Neuron | STG NeuN+ | this project | 200 | TruSeq Nano | 2x150bp | Discovery |
| 11004_Glia | STG NeuN- | this project | 200 | TruSeq Nano | 2x150bp | Discovery |
| 12004_Fib | Fibroblast | this project | 200 | TruSeq Nano | 2x150bp | Discovery |
| 12004_Neuron | STG NeuN+ | this project | 200 | TruSeq Nano | 2x150bp | Discovery |
| 12004_Glia | STG NeuN- | this project | 200 | TruSeq Nano | 2x150bp | Discovery |
| F1_Heart | Heart | this project | 200 | TruSeq Nano | 2x150bp | Discovery |
| F1_Neuron | Cortical Neuron | this project | 200 | TruSeq Nano | 2x150bp | Discovery |
| F1_Glia | Cortical Astrocyte | this project | 200 | TruSeq Nano | 2x150bp | Discovery |
| TITR | Mixture of 6 DNA | this project | 200 | TruSeq Nano | 2x150bp | Discovery |
| CONT | NA12878 | this project | 200 | TruSeq Nano | 2x150bp | Discovery |
| A1S_Heart | Heart | this project | 30 | PCR-free | 2x150bp | PCR-free vs. Nano |
| A1S_Neuron | STG NeuN+ | this project | 30 | PCR-free | 2x150bp | PCR-free vs. Nano |
| NA12889 | lymphoblastoid | Platinum | 50 | PCR-free | 2x101bp | *True/false* MEI |
| NA12890 | lymphoblastoid | Platinum | 50 | PCR-free | 2x101bp | *True/false* MEI |
| NA12891 | lymphoblastoid | Platinum | 50 | PCR-free | 2x101bp | *True/false* MEI |
| NA12892 | lymphoblastoid | Platinum | 50 | PCR-free | 2x101bp | *True/false* MEI |
| NA12877 | lymphoblastoid | Platinum | 50 | PCR-free | 2x101bp | *True/false* MEI |
| NA12878 | lymphoblastoid | Platinum | 50 | PCR-free | 2x101bp | *True/false* MEI |
| NA12879 | lymphoblastoid | Platinum | 50 | PCR-free | 2x101bp | RetroSom training |
| NA12880 | lymphoblastoid | Platinum | 50 | PCR-free | 2x101bp | RetroSom training |
| NA12881 | lymphoblastoid | Platinum | 50 | PCR-free | 2x101bp | RetroSom training |
| NA12882 | lymphoblastoid | Platinum | 50 | PCR-free | 2x101bp | RetroSom training |
| NA12883 | lymphoblastoid | Platinum | 50 | PCR-free | 2x101bp | RetroSom training |
| NA12884 | lymphoblastoid | Platinum | 50 | PCR-free | 2x101bp | RetroSom training |
| NA12885 | lymphoblastoid | Platinum | 50 | PCR-free | 2x101bp | RetroSom training |
| NA12886 | lymphoblastoid | Platinum | 50 | PCR-free | 2x101bp | RetroSom training |
| NA12887 | lymphoblastoid | Platinum | 50 | PCR-free | 2x101bp | RetroSom training |
| NA12888 | lymphoblastoid | Platinum | 50 | PCR-free | 2x101bp | RetroSom training |
| NA12893 | lymphoblastoid | Platinum | 50 | PCR-free | 2x101bp | RetroSom training |
| NA12877 | lymphoblastoid | Platinum | 200 | PCR-free | 2x101bp | RetroSom testing |
| NA12878 | lymphoblastoid | Platinum | 200 | PCR-free | 2x101bp | RetroSom testing |
| NA19238 | lymphoblastoid | HGSV* | >30 | PCR-free | 2x101bp | RetroSom testing |
| NA19239 | lymphoblastoid | HGSV | >30 | PCR-free | 2x101bp | RetroSom testing |
| NA19240 | lymphoblastoid | HGSV | >30 | PCR-free | 2x101bp | RetroSom testing |
| HG00731 | lymphoblastoid | HGSV | >30 | PCR-free | 2x101bp | RetroSom testing |
| HG00732 | lymphoblastoid | HGSV | >30 | PCR-free | 2x101bp | RetroSom testing |
| HG00733 | lymphoblastoid | HGSV | >30 | PCR-free | 2x101bp | RetroSom testing |
| HG00512 | lymphoblastoid | HGSV | >30 | PCR-free | 2x101bp | RetroSom testing |
| HG00513 | lymphoblastoid | HGSV | >30 | PCR-free | 2x101bp | RetroSom testing |
| HG00514 | lymphoblastoid | HGSV | >30 | PCR-free | 2x101bp | RetroSom testing |
| 316 WGA | 5 single clones | Bae et. al. (2018) | 5 x 30 | PCR-free | 2x150bp | RetroSom testing |
| 316 noWGA | 10 single clones/bulk | Bae et. al. (2018) | 10 x 30 | PCR-free | 2x150bp | RetroSom testing |
| 320 | 53 single clones/bulk | Bae et. al. (2018) | 53 x 30 | PCR-free | 2x150bp | RetroSom testing |
| BSM brain | cortical tissue | BSMN** | 200 | PCR-free | 2x150bp | RetroSom testing |
| BSM fib | fibroblast | BSMN | 200 | PCR-free | 2x150bp | RetroSom testing |

* Human Genome Structural Variation Consortium

** Brain Somatic Mosaicism Network Consortium

Supplementary Table 3. Sequencing features selected for RetroSom modeling.

*See additional .xlsx file.*

Supplementary Table 4. Putative somatic MEIs and post-processing.

*See additional .xlsx files.*

Supplementary Table 5. Sequences of PCR primers and probes.

*See additional .xlsx file.*

Supplementary Table 6. Evidence for ENSR00000032826.

*See additional .xlsx file.*

Supplementary Table 7. GWAS significant SNPs in the LD block around L1#1.

*See additional .xlsx file.*

SUPPLEMENTARY TEXT

Note 1. Testing RetroSom in two clone sequencing datasets

We evaluated RetroSom in two public clone sequencing datasets, 316 and 320, created by culturing individual neural cells from fetal brains and sequencing genomic DNA from each clone: dataset 316 includes 13 clones, 5 using whole genome amplification (WGA), and bulk brain and non-brain tissue; dataset 320 contains 50 clones and bulk DNA from two brain regions and one non-brain tissue^1^. In addition to being single-cell clones, these datasets differed from the Platinum dataset in sequencing method (150bp reads vs. Platinum’s 101bp reads); use of WGA in 5 of the clones for 316 (analyzed separately); and lack of family data to define true MEIs. *True* MEIs in clonal data were defined as those supported in most clones (>4 supporting reads in >80% of clones) and *false* MEIs as insertions with <3 supporting reads in >80% clones. MEIs that have many supporting reads in individual clones but are missing in others could be *true* *de novo* insertions, and thus were excluded from both the *true* and *false* groups.

Using ≥1 supporting read as the cutoff, the precision average across clones in 316-WGA, 316-noWGA, and 320 was: 52.40%, 97.03%, and 97.35%, respectively (Fig. 2A). Despite the high precision in 316-noWGA and 320, we classified an average of over 30 false positive supporting reads in each clone. Thus, we increased the stringency by only including insertions with ≥2 supporting SR and/or PE reads and improved the precision to 98.41%, 99.97%, and 99.97%, respectively, with sensitivities of 57.87%, 55.19%, and 43.96%, respectively. Using the same cutoff, the average precision for *Alu* calls were 99.76%, 99.99%, and 99.99%, respectively, with sensitivities of 83.12%, 84.48%, and 80.56%, respectively (Extended Data Fig. 2A). The number of false positives in non-WGA clones decreased to an average of 1-2 per clone, which makes it feasible to carry out PCR-based validation of all somatic MEI calls. Thus, we chose a 2-read cutoff to maintain high precision with adequate sensitivity. Our results on WGA clones suggest that amplification generates chimera artifacts that can be mistaken as somatic MEIs, and a higher stringency for the number of supporting reads is required to remove the false insertions.

Note 2. Identification of a somatic L1 insertion (L1#3) in clone sequencing dataset 320

We used RetroSom in clone sequencing datasets 316 and 320 to search for somatic MEIs in individual clones. Because each dataset consists of multiple clones, each from a single cell, we expect a somatic MEI to be heterozygous in one or more clones but be absent in most clones. Therefore, we called somatic MEIs if they were detected in < 5 clones with >5 supporting reads. We identified one L1 insertion (L1#3) in 320 basal ganglia clone 16 (320-BGclone16), supported by 17 reads and not present in any of the other 320 clones or bulk tissues (Extended Data Fig. 3). To validate this insertion, we (1) amplified both upstream and downstream junctions in one PCR product with two independent primer sets; (2) confirmed that the insertion is not found in 4 other single clones or 6 different tissues; (3) confirmed, by Sanger sequencing, that a 760bp L1 sequence (base 4536 to 5295) was inserted at chr4:56,433,538 (human hg38 coordinates), with an atypical endonuclease cutting site 5’-AATT/AT-3’; (4) confirmed a high homology (99.34%) between the inserted L1 sequence and the L1Hs consensus; (5) confirmed a 3-base target site duplication (TSD); and (6) quantified, by ddPCR, that the upstream junction is present in 22.5%, and the downstream junction is present in 24.2% of the 320-BGclone16 cells. We could not detect this insertion in 4 other clones: frontal lobe (FR) clone 3, FR clone 6, basal ganglia (BG) clone 12, and BG clone 25 and 6 bulk tissues: BG ventricular zone/subventricular zone (BG VZ/SVZ), BG cortex (BG CX), FR VZ/SVZ, FR CX, occipital cortex, and spleen. The truncation of the 3’-end, short TSD, and the atypical endonuclease cutting site suggest that L1#3 was likely created in an endonuclease independent process^2^. The absence in primary brain tissue, as well as the low mosaicism (<50%) in a single clone, both imply that it is either very rare in the brain or happened relatively early in the neural cell culture *in vitro*.

Note 3. Testing RetroSom in PCR-free and PCR-based libraries

We re-sequenced two specimens, A1S heart and A1S NeuN+, to 30x-coverage, using PCR-free sequencing libraries and 1 μg of genomic DNA each, and compared the MEI calling accuracy to two sets of six PCR-based (TruSeq Nano, ~10 PCR cycles) datasets created from the same tissues (Extended Data Fig. 4A). The *true* and *false* MEIs of A1S were selected based on their presence in all 20 libraries, including 18 TruSeq Nano (3 cell fractions) and 2 PCR-free sequencing datasets. *True* MEIs were selected as the insertions that were highly supported in most of the libraries (>4 supporting reads in >80% libraries), while *false* MEIs were selected as the insertions that were missing or poorly supported in most of the libraries (<3 supporting reads in >80% libraries).

TruSeq Nano data contained more sequencing errors, especially around low complexity regions, likely due to PCR polymerase slippage (Extended Data Fig. 4B**)**^3^. Errors made around reference poly(A) and poly(T) sites, in particular, can be mistaken as novel MEIs because they disrupt the alignment to the human reference genome but can be mapped to 3’ poly(A) tails in mobile element consensus sequences. As expected, significantly more false supporting reads were detected using TruSeq Nano than using PCR-free libraries: A1S heart L1 by 3.2 fold (95% CI: 2.6-3.9), A1S heart *Alu* by 1.4 fold (95% CI: 1.2-1.7), A1S NeuN+ L1 by 11.6 fold (95% CI: 7.8-15.4), and A1S NeuN+ *Alu* by 2.6 fold (95% CI: 2.1-3.1). RetroSom successfully removed these false MEIs by selecting against junctions of inactive transposons, improving filtering of PCR duplicates, and removing low-complexity reads.

1.	Bae, T. *et al.* Different mutational rates and mechanisms in human cells at pregastrulation and neurogenesis. *Science* **359**, 550–555 (2018).

2.	Morrish, T. A. *et al.* DNA repair mediated by endonuclease-independent LINE-1 retrotransposition. *Nat. Genet.* **31**, 159–165 (2002).

3.	Viguera, E., Canceill, D. & Ehrlich, S. D. Replication slippage involves DNA polymerase pausing and dissociation. *EMBO J.* **20**, 2587–2595 (2001).

**Acknowledgements**

We thank Wing H. Wong and Jiang Chao from Stanford University, Alexander Z. Wang and Nadya Bosch for constructive comments on the manuscript. We thank Joel E. Kleinman, Thomas H. Hyde and Daniel R. Weinberger from Liber Institute for Brain Development for providing the BSMN common brain tissue, and Liana Fasching from Yale University for extracting the BSMN common brain DNA. This work utilized computing resources provided by the Stanford Genetics Bioinformatics Service Center. **Funding:** This work was supported by Eureka Grant R01MH094740 from National Institute of Mental Health, and by the Stanford Schizophrenia Genetics Research Fund. The mixing-genome DNA sequencing and BSMN common brain sequencing data were generated as part of the Brain Somatic Mosaicism Network (BSMN) Consortium, supported by: U01MH106874, U01MH106876, U01MG106882, U01MH106883, U01MH106883, U01MH106884, U01MH106891, U01MH106891, U01MH106891, U01MH106892, U01MH106893, U01MH108898 awarded to: Nenad Sestan (Yale University), Flora Vaccarino (Yale University), Fred Gage (Salk Institute for Biological Studies), Christopher Walsh (Boston Children’s Hospital), Peter J. Park (Harvard University), Jonathan Pevsner (Kennedy Krieger Institute), Andrew Chess (Icahn School of Medicine at Mount Sinai), John V. Moran (University of Michigan), Daniel Weinberger (Lieber Institute for Brain Development), and Joseph Gleeson (University of California, San Diego). B.Z. is funded by NHLBI Grant T32 HL110952. A.E.U. is a Tashia and John Morgridge Faculty Fellow of the Stanford Child Health Research Institute. Flow cytometry sorting was performed on an instrument in the Stanford shared FACS facility obtained under NIH S10 Shared Instrument Grant (S10RR025518-01).

Members of The Brain Somatic Mosaicism Network

**Boston Children's Hospital**: Christopher Walsh, Javier Ganz, Mollie Woodworth, Pengpeng Li, Rachel Rodin, Robert Hill, Sara Bizzotto, Zinan Zhou

**Harvard University**: Alice Lee, Alissa D'Gama, Alon Galor, Craig Bohrson, Daniel Kwon, Doga Gulhan, Elaine Lim, Isidro Cortes, Joe Luquette, Maxwell Sherman, Michael Coulter, Michael Lodato, Peter Park, Rebeca Monroy, Sonia Kim, Yanmei Dou

**Icahn School of Medicine at Mt. Sinai**: Andrew Chess, Attila Jones, Chaggai Rosenbluh, Schahram Akbarian

**Kennedy Krieger Institute**: Ben Langmead, Jeremy Thorpe, Jonathan Pevsner, Rob Scharpf,

Sean Cho

**Lieber Institute for Brain Development**: Andrew Jaffe, Apua Paquola, Daniel Weinberger,

Jennifer Erwin, Jooheon Shin, Richard Straub, Rujuta Narurkar

**Mayo Clinic**: Alexej Abyzov, Taejeong Bae

**NIMH**: Anjene Addington, David Panchision, Doug Meinecke, Geetha Senthil, Lora Bingaman,

Tara Dutka, Thomas Lehner

**Rockefeller University**: Laura Saucedo-Cuevas, Tara Conniff

**Sage Bionetworks**: Kenneth Daily, Mette Peters

**Salk Institute for Biological Studies**: Fred Gage, Meiyan Wang, Patrick Reed, Sara Linker, Ani Sarkar

**Stanford University**: Alex Urban, Bo Zhou, Xiaowei Zhu

**Universitat Pompeu Fabra**: Aitor Serres, David Juan, Inna Povolotskaya, Irene Lobon,

Manuel Solis, Raquel Garcia, Tomas Marques-Bonet

**University of California, Los Angeles**: Gary Mathern

**University of California, San Diego**: Eric Courchesne, Jing Gu, Joseph Gleeson, Laurel Ball,

Renee George, Tiziano Pramparo

**University of Michigan**: Diane A. Flasch, Trenton J. Frisbie, Jeffrey M. Kidd, Mandy M. Lam, John B. Moldovan, John V. Moran, Kenneth Y. Kwan, Ryan E. Mills, Sarah Emery, Weichen Zhou, Yifan Wang

**University of Virginia**: Aakrosh Ratan, Mike J. McConnell

**Yale University**: Flora Vaccarino, Liana Fasching, Simone Tomasi, Nenad Sestan, Sirisha Pochareddy
